# Computational Scoring and Experimental Evaluation of Enzymes Generated by Neural Networks

**DOI:** 10.1101/2023.03.04.531015

**Authors:** Sean R. Johnson, Xiaozhi Fu, Sandra Viknander, Clara Goldin, Sarah Monaco, Aleksej Zelezniak, Kevin K. Yang

**Author notes:** These authors contributed equally (Sean R. Johnson, Xiaozhi Fu).

## Abstract

In recent years, generative protein sequence models have been developed to sample novel sequences. However, predicting whether generated proteins will fold and function remains challenging. We evaluate computational metrics to assess the quality of enzyme sequences produced by three contrasting generative models: ancestral sequence reconstruction, a generative adversarial network, and a protein language model. Focusing on two enzyme families, we expressed and purified over 440 natural and generated sequences with 70-90% identity to the most similar natural sequences to benchmark computational metrics for predicting *in vitro* enzyme activity. Over three rounds of experiments, we developed a computational filter that improved experimental success rates by 44-100%. Surprisingly, neither sequence identity to natural sequences nor AlphaFold2 residue-confidence scores were predictive of enzyme activity. The proposed metrics and models will drive protein engineering research by serving as a benchmark for generative protein sequence models and helping to select active variants to test experimentally.

## Main

Nature has provided us with a wealth of proteins traditionally utilized as biocatalysts to produce various valuable natural products, from everyday commodity chemicals to life-saving pharmaceuticals^1^. Advances in DNA synthesis and recombinant DNA techniques have allowed us to clone genes that encode valuable proteins into industrial organisms such as *Escherichia coli*^2^. As a result, the usage of proteins found in nature for industrial and therapeutic purposes has been a highly successful endeavor; however, the requirements of human applications are often not satisfied by natural proteins, requiring engineering to adapt them for human needs^3^.

A traditional method for moving beyond natural proteins’ sequence and function space is to use directed evolution by starting from a natural protein and iteratively making a small number of mutations until the protein acquires the desired properties^3,4^. Unfortunately, due to the vast size of protein space, many experimental efforts may result in the characterization of non-functional proteins^5,6^, i.e. it has been shown that up to 70% of random single amino-acid substitutions result in decreased activity, with 50% being detrimental to function^7–10^. Instead, computational sequence models enable the generation of new and diverse functional examples of a protein family, thus speeding up the engineering process by uncovering previously untapped functional sequence diversity and reducing the number of non-functional sequences that need to be tested^11^. Typically, these generative protein sequence models are trained either on large collections of protein sequences, e.g. the entire UniProt database of millions of sequences^12–14^, or on a set of proteins from a particular family ^15,16^, with an ultimate goal to learn the distribution of sequences within the training set in order to sample novel sequences with generalized properties of the distribution. The underlying assumption of generative protein models is that natural proteins are under evolutionary pressure to be functional, so novel sequences drawn from the same distribution will also be functional^17^. Multiple different generative protein models have been proposed, including methods based on deep neural networks such as generative adversarial networks (GANs)^15^, variational auto-encoders (VAEs)^16,18^, and language models^13,14,19–22^, and other neural networks^23,24^, as well as statistical methods such as ancestral sequence reconstruction (ASR^)25,26^ and direct coupling analysis (DCA)^27–29^. However, comparing the performance of these methods, i.e. whether computationally generated proteins are functional, remains a challenge because of limited experimental evidence evaluating model performance, likewise, there is no experimental validation supporting computational metrics.

Typically, generative protein sequence models are evaluated by comparing distributions of generated sequences to natural controls using alignment-derived scores, e.g. identity to closest natural sequence^13,15^. The few groups that have reported results from biological assays^13,18,22,27,30,31^ have used different experimental systems, making the results difficult to compare. Moreover, experimentally comparing different systems is challenging as there are many possible factors contributing to poor expression and functionality (Supplementary Table 1), ranging from mutations disrupting protein folding and stability^32^ to the codon compositions of coding regions affecting expression levels^33,34^. Thus, computational metrics for predicting the activity of novel sequences should account for as many of these factors as possible. For instance, alignment-based metrics, such as sequence identity or BLOSUM62 scores^35^, rely on homology to natural sequences and are good at detecting general sequence properties, e.g. domains. These, however, are likely to suffer at identifying crucial functional mutations as they do not account for epistatic interactions and give equal weight to all positions rather than weighting according to evolutionary conservation at each position^36^. On the other hand, alignment-free methods do not require homology search, are fast to compute and potentially can identify all common sequence artifacts based on the residue likelihoods computed by protein language models^37^. It has been demonstrated that protein language models are sensitive to identifying pathogenic missense mutations^38^, predicting protein evolutionary velocity^39^, and even capturing viral immune-escape mutations^40^. Structure-supported metrics, including Rosetta-based scores^41^, AlphaFold2^42^ residue-confidence scores, and likelihoods computed by neural network inverse folding models^43–45^, take into account protein atom coordinates potentially directly capturing protein functionality, however, they can be impractical to compute, especially when evaluating thousands of novel sequences. Although it is important to consider various rationally chosen factors when evaluating the quality of sequences through computational metrics, it is nevertheless crucial to experimentally validate these metrics for their ability to predict the activity of novel sequences accurately.

We experimentally evaluated *in silico* metrics for their ability to predict *in vitro* enzyme activity using sequences produced by three generative protein models trained on two enzyme families, malate dehydrogenase (MDH) and copper superoxide dismutase (CuSOD). Over three rounds of experiments, including naive generation, calibration, and prospective validation steps (Fig. 1a), we developed and experimentally validated COmposite Metrics for Protein Sequence Selection (COMPSS), allowing selecting successfully up to 100% of phylogenetically diverse functional sequences. COMPSS is generalizable to any protein family, and we develop a pipeline with Colab examples for community usage. This work demonstrates a composite computational metric for evaluating generated sequences that predicts experimental success. In addition to helping select active sequences for experimental validation, the proposed metrics are a first step towards establishing a standard for evaluating the performance of current and future protein sequence models that will hopefully be a catalyst for driving progress in protein engineering.

**Figure 1.**
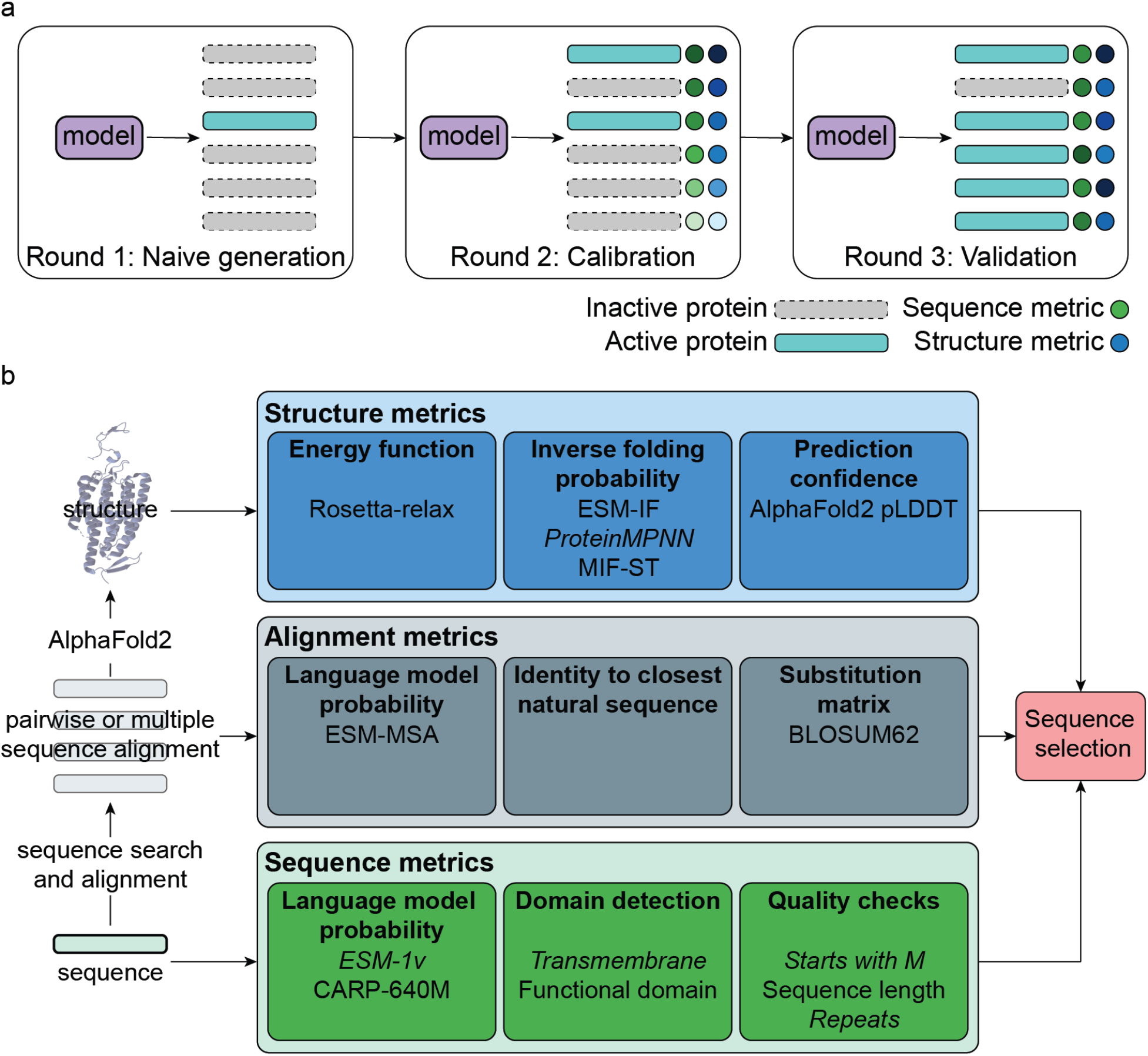
Study design. **a)** COMPSS was developed and tested over three rounds of experiments: Naive generation, calibration, and validation. **b)** COMPSS selects sequences using metrics calculated from single sequences, alignments, and predicted structures. Metrics in italics were used in the final COMPSS filter.

## Results

In this work, we consider a successful experimental validation as a protein which can be expressed and folded in *E. coli* and, after purification, have detectable activity significantly above the background in an *in vitro* assay (Methods). We focused on testing three distinct protein sequence generative models, namely, a transformer-based protein multiple sequence alignment language model ESM-MSA^46^, a convolutional neural network with attention trained as a generative adversarial network (GAN/ProteinGAN)^15^, and a phylogeny-based statistical model for the reconstruction of ancestral sequences (ASR)^26^. The latter is not a truly generative sequence model as it is constrained within a phylogeny of trees by traversing backwards in evolution without the ability to navigate sequence space in a novel direction, but it has been successfully applied for resurrecting ancient sequences^47^ and increasing enzyme thermotolerance^48^. We selected sequence models to prioritize practical simplicity, allowing them to run on workstation GPU hardware without specialized cluster equipment. To account for biological and evolutionary variations of different methodologies, we considered alignment-based, alignment-free, and structure-based metrics (Fig. 1b).

We experimentally evaluated generative models and metrics on two distinct enzyme families, EC 1.1.1.37 and EC 1.15.1.1, corresponding to malate dehydrogenase (MDH) and superoxide dismutase (SOD) families, respectively. We focus here on the copper superoxide dismutases (CuSOD) family found across all domains of life. Both MDH and CuSOD have substantial sequence diversity, numerous members in the PDB with different folds and domains, and are of physiological significance^49,50^. They are also relatively small: 300-350 residues for MDH and 150-250 residues for CuSOD, but are complex proteins active as multimers, and their activity can be assayed by the spectrophotometric readout.

### Round 1: Naive generation results in mostly inactive sequences

We started by constructing a training set of sequences to train generative models of CuSOD and MDH. We collected a total of 6,003 CuSOD and 4,765 MDH sequences from Uniprot (Supplementary Table 2) with corresponding domains for each protein family. CuSOD training sequences had only a single Sod_Cu domain, while MDH had an Ldh_1_N followed by an Ldh_1_C domain and no other Pfam domains that generally only rarely occur in 6.3% and 1.7% of sequences in both families, respectively. Before training, sequences were truncated around the annotated domains to remove possible signal peptides, transmembrane domains, and extraneous unannotated domains. After data preprocessing and, where necessary, model training and fine-tuning, we generated a total of >30,000 sequences from the ASR, ProteinGAN and ESM-MSA models (Supplementary Table 3) and selected 144 sequences for experimental validation with 18 sequences for each model as well as a set of natural test sequences for determining control success rates. All generated sequences were selected to have 70%-80% identity to the closest training natural sequence (Supplementary Table 4).

19% of all experimentally tested sequences, including natural sequences, showed *in vitro* activity (Supplementary Table 5, Supplementary Figs. 11-16). Of the CuSOD sequences, none of the test or ESM-MSA and only two of the GAN sequences displayed activity. Of the MDH sequences, none of the GAN or ESM-MSA sequences had activity, but 6 of the test sequences did. In contrast, the ASR model generated 9 of 18 and 10 of 18 active enzymes for CuSOD and MDH, respectively.

We then investigated potential reasons for poor performance in activity, including the natural test sequences. First, we observed that test sequences with predicted signal peptides or transmembrane domains in the pre-truncation sequences (Methods) were significantly overrepresented in a non-active set (one-tailed Fisher test, *p-*value = 0.046). For CuSOD, a literature search^51,52^ combined with careful examination of the assayed sequences and the available CuSOD crystal structures (PDB: 4B3E)^53^ showed that CuSOD is active as a homodimer (or sometimes a tetramer) and that the truncations we made to the natural sequences tended to remove residues located at the dimer interface, which likely interferes with enzyme expression and activity. We thus made equivalent truncations to our positive control enzymes, human SOD^154^ (hSOD, GenBank: NP_000445.1), *Potentilla atrosanguinea* (a plant) CuSOD^55^ (paSOD, GenBank: AFN42318.1) and *E.coli* SOD^56^ (E.SOD, GeneBank: NP_416173.1), and indeed this confirmed the loss of activity for hSOD and paSOD (Supplementary Figs. 17,18). We note that the over-truncation also affected the ASR sequences, yet many were still active. We speculate this may be due to the stabilizing effect widely reported in reconstructed ancestral sequences^25,57,58^.

To further test the hypothesis that poor truncation selection was responsible for the lack of observed activity in the Round 1 CuSOD natural test sequences, we assayed an additional 16 natural SOD proteins (pre-test group). In nature, eukaryotic CuSOD proteins are typically cytosolic and therefore lack a signal peptide, whereas bacterial CuSOD proteins are typically secreted^51^. CuSOD sequences were manually selected based on the kingdom of origin, eukaryotic, viral, or bacterial, and the presence of signal peptides was predicted using Phobius^59^. Sequences with predicted signal peptides were truncated at the predicted cleavage site. Two bacterial FeSOD proteins, both lacking a predicted signal peptide, and the previously characterized *E. coli* FeSO^D56^ (sodB, GenBank: NP_416173.1, as a positive control) were also assayed. 8 out of the 14 CuSOD sequences had activity, including three of the four eukaryotic enzymes and the single viral enzyme, all of which lacked a predicted signal peptide and were expressed in their full-length form (Supplementary Fig. 1, 19, 20). In the case of MDH, poor truncation selection seemed less of an issue, as 6 out of 17 test sequences were active (Supplementary Table 5, Supplementary Figs. 11a, 12a, 13a).

### Round 2: Calibration data for COMPSS

Consolidating lessons learned from the exploration of Round 1, we retrained generative models and tested additional sequences to calibrate the computational metrics. Specifically, in this round, for the training set and natural test sequences of both families, we now used only full-length natural sequences, filtering out sequences with predicted transmembrane domains and signal peptides. For CuSOD, we further restricted the set to proteins from eukaryotic or viral sources (Supplementary Table 2). For sequence generation, we used the same method as in Round 1 for generating ASR and GAN sequences but modified the ESM-MSA sampling procedure to improve the quality of generated sequences (Supplementary Table 3). ESM-MSA sampling for Round 1 used MSAs composed of randomly selected training sequences and masked and sampled across the entire MSA. For Round 2, only one training sequence from the MSA was masked and sampled, and the MSA was composed of the training sequences most similar to the sampled sequence. We selected 18 sequences each from the ASR, GAN, and ESM-MSA. Sequences generated with the revised ESM-MSA sampling method tended to have higher scores on metrics, including ESM-1v and identity to the closest training sequence (Supplementary Fig. 2). Only 13 test natural sequences were selected, as we had already screened five similar natural sequences in the remediation for Round 1. The number of expressed enzymes with activity above the background was substantially higher than in Round 1, with 66% of natural controls showing activity when expressed in *E. coli* and at least 50% of active synthetic sequences for every model/enzyme-family combination except for GAN/MDH, where only two out of 18 sequences were active (Supplementary Table 5, Supplementary Figs. 21-24).

To calibrate the metrics against enzymatic activity for each experimentally tested sequence, we generated alignment-based (Identity, BLOSUM62^35^, ESM-MSA) and sequence-only alignment-free (CARP-640M^60^, ESM-v1^37^) metrics. We also predicted AlphaFold2 structures^42,61^ for all of the selected sequences and calculated structure-based metrics using Rosetta^41^ and metrics based on the inverse folding neural networks ProteinMPNN^45^, ESM-IF^43^, and MIF-ST^44^. Besides the identity range, the experimentally tested sequences were selected to span the entire range of scores on each metric (Supplementary Table 4). Apart from alignment-based ESM-MSA, none of the metrics substantially correlated with sequence identity, suggesting that our chosen metrics contain orthogonal information that is not accounted for entirely by sequence identity (Supplementary Fig. 4). On the other hand, structure-based metrics overall displayed substantial correlations with each other, with the highest correlations observed between inverse folding neural network scores (Fig. 2b, Supplementary Fig. 3).

**Figure 2.**
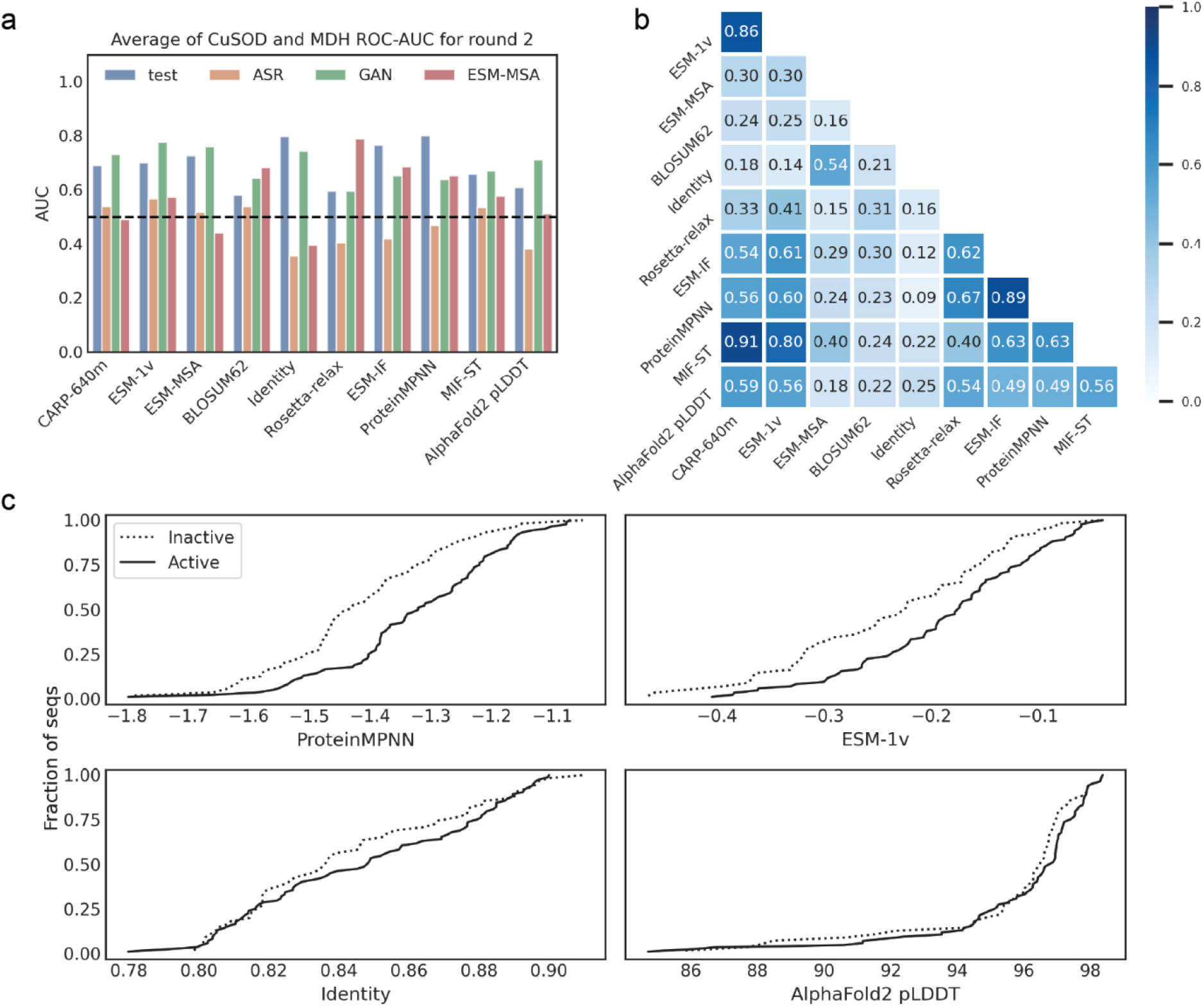
Computational metrics of sequences experimentally tested in Round 2. **a)** Spearman correlations between metrics Average of CuSOD and MDH. **b)** AUC-ROC scores of activity vs metrics. Average of CuSOD and MDH. **c)** Empirical cumulative distribution function curves of active (solid lines) and inactive (dotted lines). Curves are for the pooled results of all generative models and both enzyme families.

AUC-ROC curve scores between activity and each metric (Fig. 2a, Table 1, Supplementary Figs. 5, 6) indicate that they could help predict activity in novel sequences, but generally, none stands out superior to the others. Structure-based metrics, on average, best explain the enzymatic activity, showing 0.67 AUC-ROC among all models and enzyme families. In contrast, we did not observe differences in AlphaFold2 residue-confidence scores pLDDT scores between active and inactive generated enzymes in (Wilcoxon rank-sum test *p*-value = 0.31): generated sequences result in high-quality AlphaFold2 structures irrespective of their activity (Fig. 2c). Likewise, sequence identity did not explain enzymatic activity (Table 1, Fig. 2c).

**Table 1.**
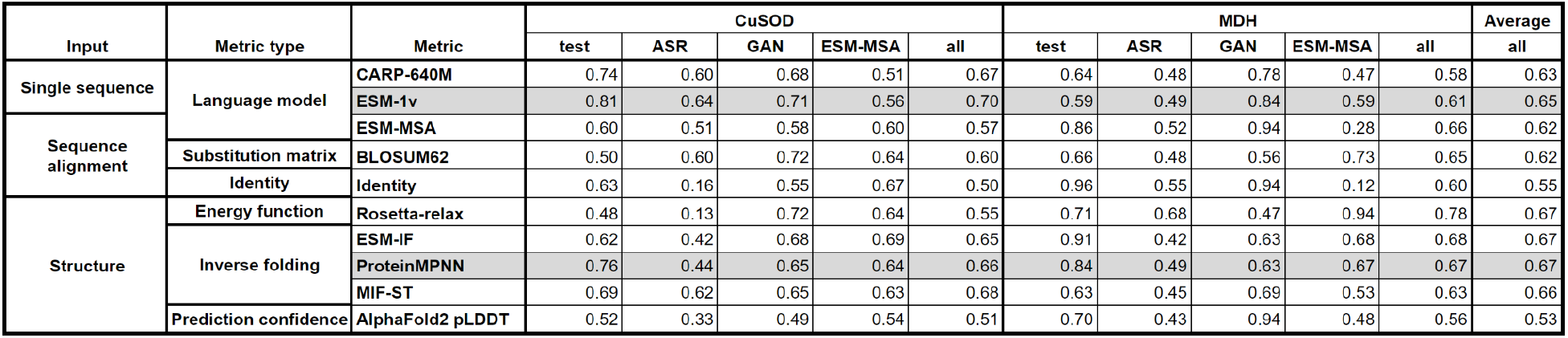
AUC-ROCs of each metric vs experimentally measured activity in Round 2. The ESM-1v and ProteinMPNN metrics are highlighted to indicate that they were chosen to be part of the filter for Round 3.

### Round 3, validation: COMPSS metric enriches active protein generation

Next, we aimed to devise a sequence prioritization technique or *in silico* filter that would enable us to perform virtual screening of large numbers of generated sequences to identify probable active sequences with an identity <80% to the closest natural sequence. Based on Round 2 (Fig. 2, Table 1), no single metric would be sufficiently generalisable to screen against multiple sequence failure modes (Supplementary Table 1), so we tested a filter composed of a combination of metrics. To combine different data types, we used the ESM-1v and ProteinMPNN metrics (Fig. 3a, Supplementary Fig. 9). These metrics are attractive because ESM-1v is sequence-based and ProteinMPNN considers structural information, neither of which is strongly correlated with sequence identity (Fig. 2a). Although most structure-based metrics performed similarly and could be interchanged (Table 1), ProteinMPNN was deemed the most practical due to its lower parameter count. Rosetta-relax performed best on MDH sequences but is computationally intensive compared to other structure-based metrics. The ESM-1v protein language sequence model had the best performance for alignment-free metrics, with an AUC-ROC up to 0.81 for ProteinGAN MDH sequences and fast computation. Furthermore, the correlation between the scores of the two metrics was only moderate, with an average Spearman’s ρ = 0.60 (Fig. 2a), suggesting that the metrics may capture distinct features.

**Figure 3.**
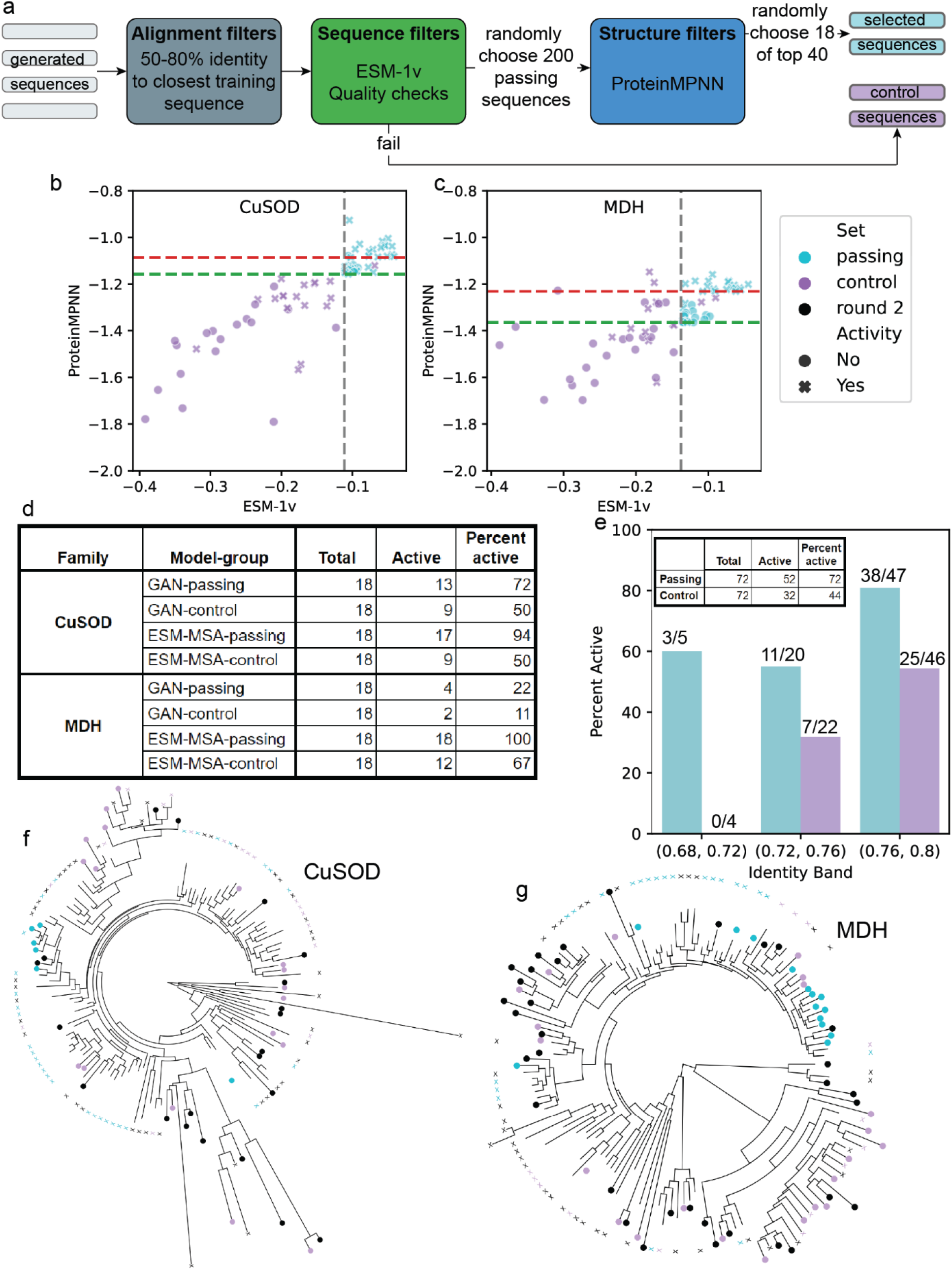
Round 3 selection and experimental results. **a)** Computational filter for selecting sequences to screen in Round 3. **b)** CuSOD and **c)** MDH ESM-1v and ProteinMPNN scores for Round 3 selected passing (teal) and control (violet) sequences. The vertical dashed grey line indicates the top 10 percentile ESM-1v score cutoff calculated from the test sequences. Horizontal dashed lines are the ProteinMPNN scores of the 40th-ranked sequences of the 200 for which the scores were calculated for each model-family combination, ProteinGAN (lower, green), ESM-MSA (upper, red). Control sequences to the right of the grey line are possible if they fail one of the quality checks. **d)** Tabulation of active enzymes in each group. **e)** Active enzymes by identity band. **f)** Phylogenetic tree of CuSOD enzymes screened in Round 2 (black) and Round 3 (passing, teal; control, violet). X indicates an active sequence, filled circle indicates an inactive sequence. **g)** Phylogenetic tree of MDH sequences.

To select a threshold for ESM-1v scores that would enrich active proteins for any new enzyme family, we analyzed the experimental data of natural test sequences from Round 2. We found that the highest enrichment of active sequences occurred at about the 20th percentile of ESM-1v scores, with the top 23% and 17% of CuSOD and MDH scores, respectively, having the highest fraction of active proteins (Supplementary Fig. 8). For prospective validation of our sequence prioritization strategy, in Round 3, we used the 10th percentile cut-off from the natural sequences, making the threshold more stringent, as, in practice, the score should be derived from untested natural sequences. To validate our strategy, we focused on GAN and ESM-MSA models, as ASR sequences performed consistently well in our Rounds 1 and 2 and in numerous literature report^s25^ and would therefore be a poor test for our filter. After performing quality checks for sequences starting with methionine, lacking long repeats, and transmembrane domains, we randomly selected 200 ESM-1v threshold passing sequences within the 50%-80% target identity band to natural sequences for each model and enzyme family. Then for each selected sequence, we predicted structures using AlphaFold2, calculated ProteinMPNN scores, and randomly selected 18 sequences from those with ProteinMPNN scores in the top 40 for each model and enzyme family combination. For each passing sequence selected for experimental validation, a negative control sequence was randomly chosen from the sequences failing the sequence filters. Control sequences had an identity to the closest training sequence within 1% of the passing sequence. The stringency of the ESM-1v filter led to some phylogenetic bias in the sequence selection, particularly for MDH; nevertheless, the set of screened enzymes covered approximately the same space in Round 3 as in Round 2 (Fig. 3f,g).

A total of 144 selected passing and control sequences were expressed in *E. coli*, purified, and assayed for activity in the same way as in previous rounds (Supplementary Figs. 25-30). Most of the sequences the filter selected showed *in vitro* activity, including 94% (17 out of 18) and 100% of ESM-MSA CuSOD and MDH enzymes, respectively (Fig. 3b-e, Supplementary Table 5). Furthermore, considering the sequences from both models and enzyme families, the selected sequences had a 62% significantly higher success rate (Fisher test *p*-value = 0.0012) than the sequence-filter-failing control sequences, with a total of 72% active sequences (Fig. 3e, Supplementary Fig. 10).

To further validate the COMPSS pipeline, we tested it against previously published datasets of experimentally characterized sequences from six additional families generated by models with architectures distinct from the generative models used in this work (Supplementary Table 6), i.e. synthetic enzymes from five evolutionarily distinct lysozyme families generated by the single-sequence language model ProGen^13^, and chorismate mutases generated by the DCA model bmDCA^27^. For five of the six families, there was a higher rate of functional enzymes among those passing the COMPSS sequence filter than among those failing the filter, and AUC scores for the ProteinMPNN structural metric were above 0.5 for all families, ranging from 0.6 to 1.0 (Fig. 4).

**Figure 4.**
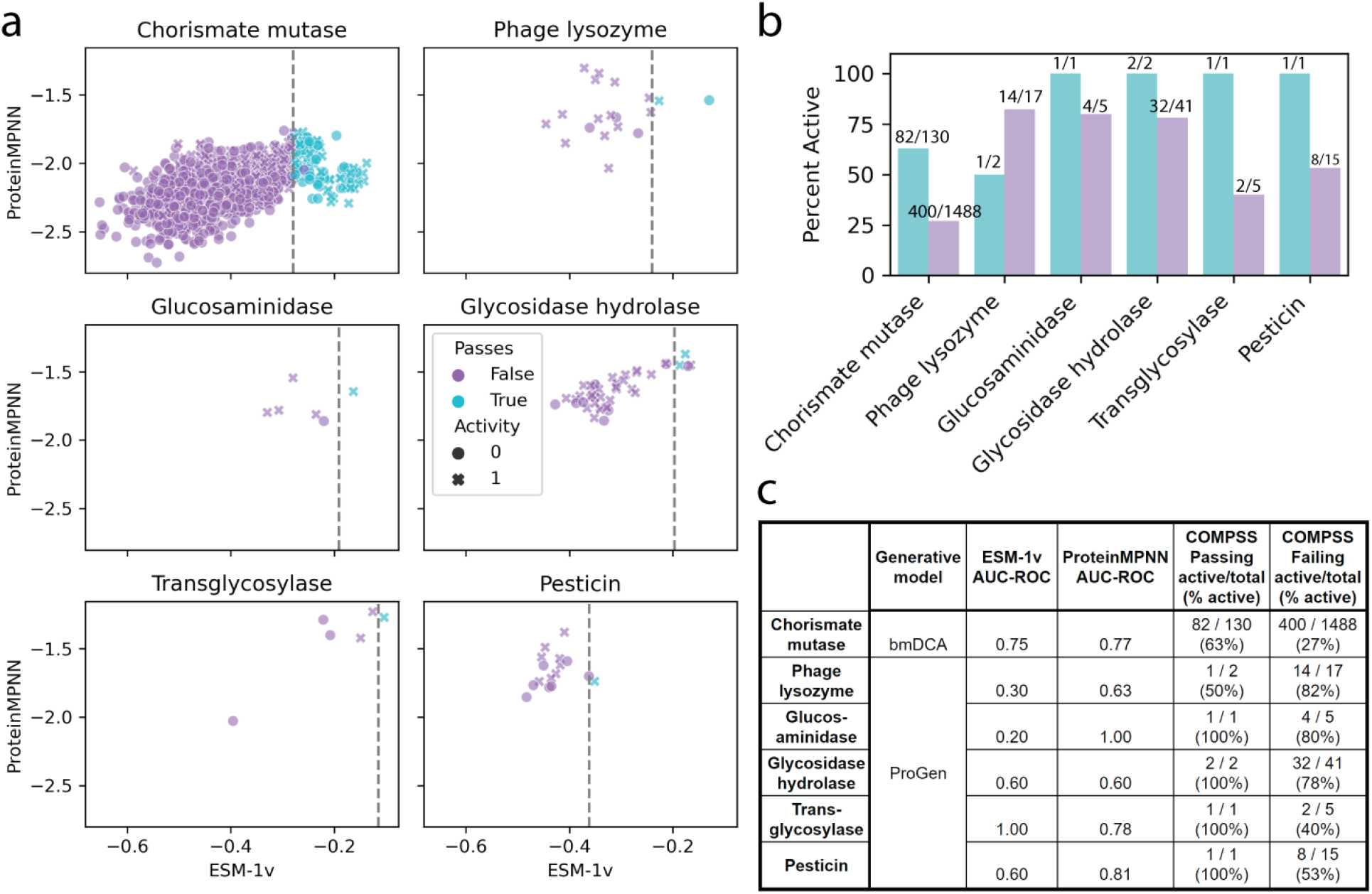
Validation of COMPSS against generated enzymes from six previously unseen protein families and two new models. **a)** ESM-1v and ProteinMPNN scores of generated enzymes from the six families. The vertical dashed grey line indicates the top 10 percentile ESM-1v score cutoff calculated from natural sequences from the same families. **b)** Proportion of active enzymes stratified by family and whether they pass the COMPSS sequence filter. **c)** Summary of COMPSS performance on the six families from publicly available data.

## Discussion

Discovering new enzyme variants is difficult because it requires understanding their molecular mechanisms, expression and folding supporting their intended activity while operating within biological and physical constraints. Generative protein sequence models mimic these constraints by learning underlying protein sequence distributions to sample novel sequence variants. With the advent of deep neural network technologies, there have been tremendous breakthroughs in generative protein designs^62^, allowing the design of *de novo* proteins with specified folds^45,63^. Due to emergent complexity defining enzymes’ catalytic functions (Supplementary Table 1), predicting which novel enzyme sequence will express and fold in a soluble and active form remains challenging, ultimately limiting the discovery of novel enzymes. To facilitate novel enzyme discovery from non-natural sequence spaces, we experimentally evaluated a diverse set of *in silico* metrics to determine their efficacy in predicting novel sequence activity. We then selected a subset of the metrics to form the core of COMPSS (COmputational Metrics for Protein Sequence Selection). We validated the filter’s effectiveness by prospectively testing synthetic selected proteins resulting in up to a two-fold increase in the number of active protein sequences (Fig. 3d). Similar results were obtained by independently validating COMPSS on previously published datasets^13,27^ of six enzyme families generated by models not considered in the present study (Fig. 4).

Our experimentally validated end-to-end framework for generating and selecting new active enzyme variants (Box 1) consists of three steps. The first is curating sequences to develop a clean training dataset. Machine learning is a data-centric practice as much as it is algorithmic; as such, dataset curation was crucial for both tested generative models, i.e. neither evolutionary scale deep transformer neural network (ESM-MSA) nor the local and much smaller ProteinGAN model performed well in the naive setting in Round 1 (Supplementary Table 5). In Round 2, we observed that removing sequences containing transmembrane domains and focusing on sequences without signal peptides enriched the dataset for natively soluble cytosolic proteins and led both neural network models to generate active enzymes at a rate of up to 67% (Supplementary Table 5). In this second round, we also focused on generating synthetic sequence diversity in order to calibrate our metrics. Specifically, to inform the design of a sequence selection strategy, we selected sequences with a broad range of scores on the metrics and observed which metrics correlated with activity (Fig. 2). For the third validation round of *in vitro* measurements, we combined sequence-based quality checks, the sequence-based ESM-1v metric, and the structure-based ProteinMPNN metric to score and select generated sequences, resulting in the enrichment of active sequences (Fig. 3a, Supplementary Fig. 9). Based on data from the two rounds of experimental measurements, we found that while AlphaFold2^42^ accurately predicts protein structures from MSAs, and AlphaFold2 embeddings generalize multiple protein properties^64^, the frequently used residue-confidence pLDDT score does not distinguish enzyme activity within sets of homologous synthetic sequences (Fig. 2a,c). AlphaFold2 produces high-confidence structures even for synthetic sequences that do not result in active catalytic proteins (Fig. 2c, Supplementary Fig. 4). Likewise, neither identity nor sequence similarity is sensitive enough to capture the functional differences between evolutionarily-related sequences (Table 1, Fig. 2a,c). On the other hand, likelihoods from protein language models and from models that predict sequence from the structure moderately explain *in vitro* enzyme activity and only weakly correlated with each other (Table 1, Fig. 2a). Therefore, we chose to use one language model (ESM-1v) and one model that predicts sequence from structure (ProteinMPNN) in the final COMPSS pipeline.

Our study is not intended to benchmark generative models against each other but instead to evaluate metrics to identify those widely applicable across generative models and enzyme families. As such, we did not do much optimization of models hyperparameters, instead focusing on learning how to identify active enzymes from pools of generated sequences. However, we do find that ancestral sequence reconstruction outperforms more complex neural network models at naive generation, indicating room for improvement in deep generative models of protein sequences. We observed a wide range of AUC scores for different combinations of a generative model, enzyme family, and metric, likely due to a combination of factors. Different models and enzyme families may be prone to different failure modes captured by different metrics. Some metrics use similar underlying models as some of the generative models, which may lead to an overfitting effect. Despite evaluating in total over 2200 enzyme variants from 8 different enzyme families, given the sheer size of protein sequence space, an even more extensive dataset would be desirable to tease apart the complex interplay between generative models, metrics, and protein families and to explain that interplay in biologically meaningful terms.

We show that by carefully curating training data for sequence generation and prioritizing sequences for experimental testing using a multipart filter, as many as 100% of enzymes having *in vitro* activities can be achieved, with sequence identities between 70% and 80% to the closest naturally occurring enzymes. To aid others in using these methods, we provide code in the form of Google Colab notebooks capable of generating new sequences using the ESM-MSA model and calculating metrics for any user-supplied protein sequences or structures. Our dataset of more than 440 experimentally tested enzymes and multiple metrics can serve as reference benchmarks for further improved metrics for predicting the function of generated sequences. The presented end-to-end workflow provides a powerful and flexible framework for developing diverse libraries of active enzymes, enabling deeper explorations of functional sequence space.

**Box 1:**
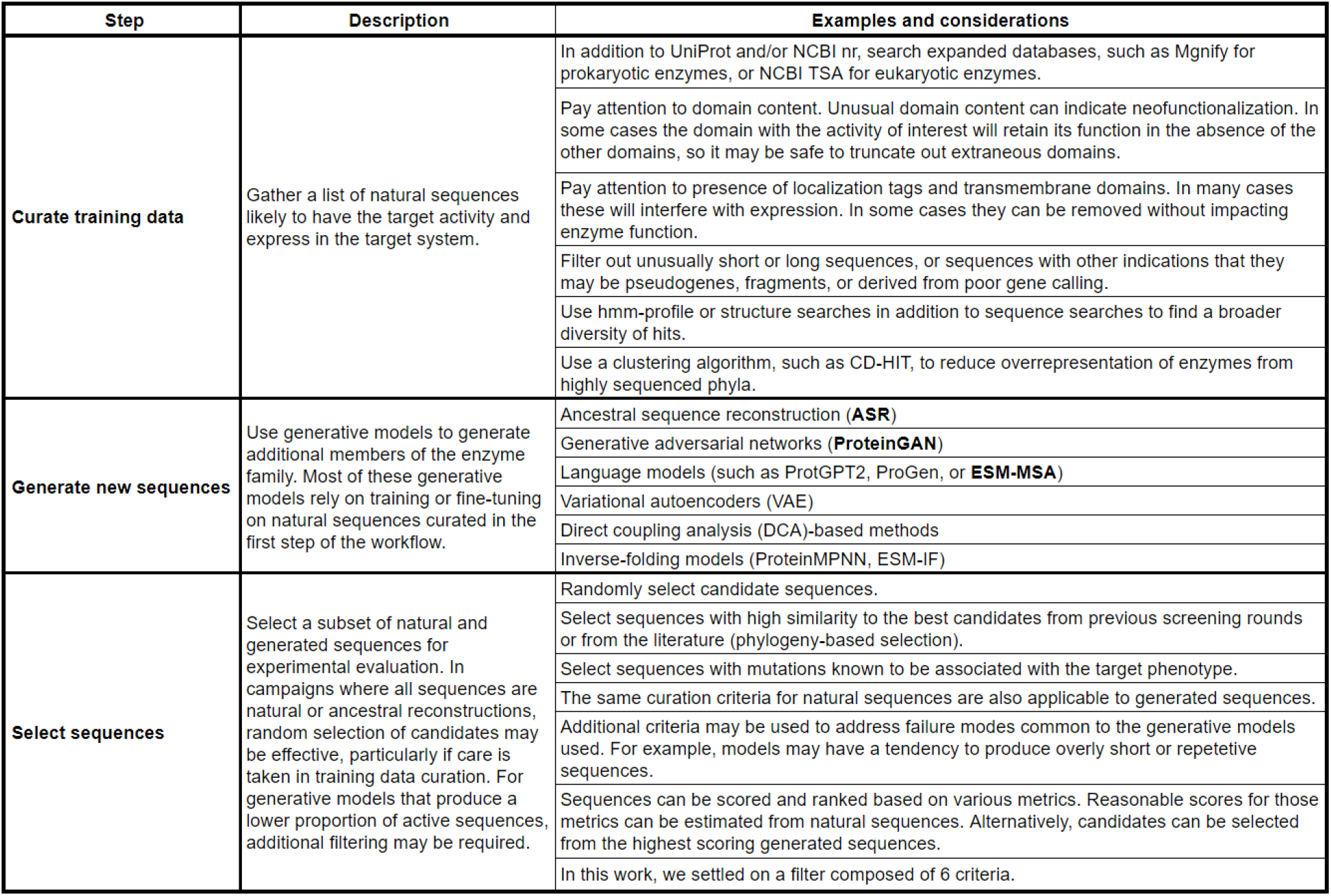
Framework for generating and selecting new active enzyme variants.

## Data and Code Availability

All generated sequences, curated natural sequences, train/test splits, predicted structures, metrics scores, phylogenetic trees, and tabulations of experimental results are available at a Zenodo deposit (https://doi.org/10.5281/zenodo.7688667)

Code for regenerating figures, and links to Colab notebooks for calculating metrics and generating sequences using ESM-MSA are available on Github (https://github.com/seanrjohnson/protein_scoring)

Locally executable code for generating sequences from ESM-MSA is also available on Github (https://github.com/seanrjohnson/protein_gibbs_sampler)

## Supplementary information

Methods, Supplementary Figs. 1-30, Supplementary Tables 1-7

## Acknowledgements

We gratefully thank Vladimir Potapov for reviewing a draft of this manuscript. We thank Philip Rosenfield and Irene Money at Microsoft Research New England for assistance in securing internal funding for wet-lab experiments and Nicolo Fusi for useful discussions. We thank the Chalmers Center for Computational Science and Engineering (C3SE) and the Swedish National Infrastructure for Computing (SNIC) for providing computational resources. Computing resources at the Chalmers Center for Computational Science and Engineering (C3SE) were partially funded by the Swedish Research Council through grant agreement no. 2018-05973. Mikael Öhman and Thomas Svedberg at C3SE are acknowledged for technical assistance. The study was supported by SciLifeLab funding (A.Z.), Swedish Research council (Vetenskapsrådet) starting grant no. 2019-05356 (A.Z.), Formas early-career research grant 2019-01403 (A.Z.) and Marius Jakulis Jason foundation (A.Z.).

