## Supplementary_methods_figures_and_tables for "Computational Scoring and Experimental Evaluation of Enzymes Generated by Neural Networks"

#### Data curation

##### Round 1 CuSOD

Using the InterPro[1] web interface, all UniProt[2] sequences containing exactly one Sod_Cu Pfam[3] domain, and no other Pfam domains, were downloaded. Hmmsearch (from the Hmmer3 suite)[4,5] was used to find the coordinates of the Sod_Cu domain envelope in each sequence. From each sequence, the region containing the envelope, plus up to 10 amino acids at the N-terminus, if present, and either 0 additional amino acids at the C-terminus (if the end of the domain envelope was less than 10 amino acids from the end of the protein), or 10 additional amino acids at the C-terminus (if at least 10 additional amino acids were present in the protein). We intended to extract up to an additional 10 amino acids on the C-terminus, as with the N-terminus, but a software bug led to some extractions ending at the end of the domain envelope.

All extracted CuSOD domain sequences containing ambiguous or non-canonical amino acids were discarded. Remaining domains with a length outside a window of 1 standard deviation from the median length were discarded. Remaining domains were clustered using CD-HIT[6] with an 80% identity threshold, and the cluster center sequences were retained. The cluster centers were randomly sorted 80%-20% into a “training” and “test” set, containing 4802 and 1201 sequences, respectively.

A compact, representative, MSA was generated by randomly selecting 1500 sequences from the training set, running MUSCLE (v3.8)[7] and then removing all sequences with an amino acid in any MSA column containing at least 99% gaps. The alignment and filtering was repeated until no sequences were removed, resulting in an MSA with 445 sequences.

##### Round 1 MDH

Using the InterPro web interface, all UniProt sequences containing exactly one Ldh_1_N Pfam domain followed by exactly one Ldh_1_C Pfam domain were downloaded. The Ldh_1_N / Ldh_1_C domains cover both lactate dehydrogenases (LDH) and MDH activities, but we were only interested in the enzymes with MDH activity.

To distinguish sequences with the two activities, we used the UniProt web interface to find reviewed LDH and MDH enzymes based on Enzyme Comission (E.C.) number[8], 1.1.1.27 for LDH, and 1.1.1.37 for MDH, from the reviewed, SwissProt, subset of UniProt. Muscle was used to make MSAs of the two enzyme groups, and hmmbuild (from the Hmmer3 suite) was used to build profile hidden markov models of both MSAs. Hmmsearch was used to score each UniProt Ldh_1_N/Ldh_1_C sequence against the MDH and LDH profiles, and those that had a stronger match to the MDH profile were retained.

Additional processing was done exactly as with the Round 1 CuSOD data curation, resulting in 3812 training and 953 test sequences, and a training MSA of 835 sequences.

#### Quantifying domain architectures

To quantify the number of proteins with just the desired number of target domains and no other domains, we used hmmsearch against UniProt, evalue cutoff 0.001, to identify all proteins with the Pfam domains of interest, Sod_Cu for CuSOD, and Ldh_1_N and Ldh_1_C for MDH. For MDH, we further filtered so that our list of proteins only included those with both Ldh_1_N and Ldh_1_C. On the hits from those searches, we ran hmmscan[5], evalue cutoff 0.001, Z 230895644 (which is the size of the UniProt database), and filtered out domain annotations that overlapped more than 60% with better scoring domain annotations. From the list of annotated sequences, we counted the number of sequences containing just the desired domain architecture, a single CuSOD annotation and no other domains, or a single Ldh_1_N followed by a single Ldh_1_C and no other annotations. For CuSOD, 1,632 out of 25,701 proteins (6.3%) had aberrant architectures. For MDH, 1,127 out of 65,639 (1.7%) had aberrant architectures.

#### Round 2 CuSOD pre-test

UniProt CuSOD proteins were obtained as described above (Round 1 CuSOD). The kingdom of origin for each sequence was obtained from the UniProt annotation. Transmembrane domains and signal peptides were predicted using Phobius[9]. Sequences with transmembrane domains were discarded. Signal peptides were removed from sequences predicted to contain them. A set of 14 representative CuSOD and two FeSOD proteins were manually selected for experimental screening, including eukaryotic, viral, and bacterial proteins predicted to not contain signal peptides, and bacterial proteins with predicted signal peptides removed.

#### Rounds 2 and 3 CuSOD

All eukaryotic transcriptomes available from the NCBI Transcriptome Shotgun Assembly (TSA) sequence database[10] were downloaded. Transdecoder (https://github.com/TransDecoder/TransDecoder) was used to extract the protein sequences from the transcriptomes. Hmmsearch[5] was used to identify proteins with exactly 1 CuSOD domain and no other Pfam domains. This set of proteins was combined with the list of eukaryotic and viral CuSOD proteins from UniProt. From the combined list, all proteins with at least 1 transmembrane domain or a signal peptide, as predicted by Phobius, were discarded. All sequences extending beyond the CuSOD domain envelope more than 25 amino acids at the N-terminus or 15 amino acids at the C-terminus were discarded. All sequences smaller than 150 amino acids or larger than 250 amino acids were discarded. All sequences with more than 85% identity, based on usearch[11] search_global, to a sequence screened in a previous round were discarded. Remaining sequences were clustered using CD-HIT at a 90% identity threshold, and the cluster centers were retained. The remaining sequences were split 90%-10% into 2543 training and 283 test sequences. A training MSA was generated as described for Round 1 CuSOD, except that a 99.5% column gap threshold was used, giving an MSA with 947 sequences.

#### Rounds 2 and 3 MDH

Hmmsearch[5] was used to identify sequences in the Mgnify[12] metagenomics database containing exactly one Ldh_1_C and one Ldh_1_N domain, and no other Pfam domains. The list of Mgnify proteins was added to the list of UniProt proteins with the same architecture (curation described above) before any additional filtering. From the combined set, sequences were thrown out if they met any of the following conditions: longer than 353 amino acids, shorter than 290 amino acids, contains ambiguous or non-canonical amino acids, does not start with M, matches the LDH SwissProt profile (described above) better than the MDH SwissProt profile. Phobius transmembrane domain and signal peptide prediction was run on the remaining sequences. All sequences containing a predicted transmembrane domain or a predicted signal peptide were discarded. All sequences with an N-terminal or C-terminal overhang going beyond 20 amino acids past the beginning of the Ldh_1_N envelope or the end of the Ldh_1_C envelope were discarded. Remaining sequences were clustered at a 90% identity threshold with CD-HIT and the cluster centers retained. All sequences with identity greater than 85%, based on usearch search_global, to a sequence experimentally screened in Round 1 were discarded. The remaining sequences were split 90%-10% into 7187 training and 799 test sequences. A training MSA was generated as described for Round 1 CuSOD, except that a 99.5% column gap threshold was used, giving an MSA with 975 sequences.

##### Phylogenetic trees

Trees were made by FastTree from MSAs generated by MAFFT[13]. Trees were rooted and the midpoint and rendered using ETE3[14].

#### Chorismate mutase and lysozymes

1130 chorismate mutase natural sequences were acquired from the supplementary materials of Russ et al. (2020)[15]. Lysozyme natural sequences were acquired by downloading the corresponding Pfam profiles (Supplementary Table 6) from the InterPro website (https://www.ebi.ac.uk/interpro/) and searching for hits in UniProt using hmmsearch[5], evalue cutoff 0.001, Z 1000. To extract the lysozyme domains from the hits, we extracted sequence regions including the domain hit envelope plus extended regions on either side. Extensions were different for each profile and were determined by examining the natural enzymes tested by Madani et al. (2023)[16]. PF00969: 60 N-term, 20 C-term; PF05838: 10 N-term, 80 C-term; PF01832: 5 N-term, 5 C-term; PF06737: 5 N-term, 10 C-term; PF16754: 60 N-term, 20 C-term. The lysozyme natural sequences were clustered using CD-HIT at a 90% identity threshold, and the cluster centers were retained. Top 10% ESM-1v thresholds were calculated from the natural sequences for each family. Generated chorismate mutase and lysozyme sequences were acquired from the supplementary materials of the corresponding papers, structures were predicted using AlphaFold2, and ProteinMPNN and ESM-1v scores and sequence quality checks were calculated in the same way as for the MDH and CuSOD generated sequences. Chorismate mutase enzymes were categorized as “active” if the “norm r.e.” score was > 0.42. Lysozyme sequences were categorized as “active” if the “functional?” column contained the value “TRUE”, and inactive if that column contained the value “FALSE”. There were 10 generated lysozymes where “functional?” was marked as “-”, we omitted these 10 from our analysis. When applying the COMPSS sequence filter to chorismate mutase and lysozyme sequences, we omitted the identity filter and the “starts with M” filter, because these filters would eliminate most sequences in the datasets.

### Generative models

#### ESM-MSA-1b sampling

Sequences were generated using iterative masking and sampling using the ESM-MSA-1b model [17]. ESM-MSA-1b is a neural network model trained to fill in the WT amino acids in masked positions of a protein multiple sequence alignment. The model can be used to generate new sequences by running the MSA masking and prediction iteratively, each time replacing the WT amino acids at the masked positions with an amino acid drawn from the probability distribution returned by the model. Using masked language models to generate new sequences was first proposed by Wang and Cho[18], and the strategy has been applied to protein sequences in at least 3 prior works[19–21].

Key parameters for sampling are the composition of the MSA, which sequences in the MSA are masked and sampled, what percent of the residues are masked in each iteration, how many burn-in iterations to run, where amino acids are drawn from the full probability distribution, and how many top-k iterations to run, where amino acids are drawn only from the k most probable amino acids.

For Round 1, for both MDH and CuSOD, the MSA size was 64 sequences. The 64 sequences were randomly drawn in pre-aligned format from the training MSA. A random 10% of amino acids from the entire MSA were masked and sampled in each iteration. 20 iterations were run, which is equivalent two complete passes over the all positions. All iterations were burn-in iterations. The process was repeated 79 times with different random draws from the training MSA, to generate a total of 5000 new sequences.

For Rounds 2 and 3, for both MDH and CuSOD, the MSA size was 32 sequences. Masking and sampling was done on just a single sequence at a time, the "template sequence". Every training sequence was used as a template sequence. The rest of the MSA was filled out by using phmmer (from the Hmmer3 suite) to search the training sequences using the template sequence as a query, and keeping the top 31 hits, not including the self-hit of the template sequence, and an MSA of the set of 32 sequences was generated using MAFFT (v7.487) [13] in globalpair mode. A random 10% of amino acids from the template sequences were masked and sampled in each iteration. 30 iterations were run, the first 20 were burn-in iterations, the final 10 had top-k of 1.

For Round 2, one new sequence (2543 CuSOD, and 7187 MDH) was generated for each training sequence. For Round 3, the list was further expanded by the same method by adding an additional seven resamplings of each CuSOD train and three resamplings of each MDH sequence, for a grand total of 28,748 MDH sequences and 20,344 CuSOD sequences.

Code for generating protein sequences by sampling from ESM-MSA can be found here:
[seanrjohnson/protein gibbs sampler: Gibbs sampling for generating protein sequences (github.com)](https://github.com/seanrjohnson/protein_gibbs_sampler)

#### ProteinGAN

Generative adversarial models were trained using the training sets for CuSOD, and MDH respectively. Then, for each family, sequences were generated for each family by sampling vectors from the latent space using a truncated normal distribution. For Rounds 1 and 2, 10,048 sequences were generated for each family. For Round 3, 560,016 and 160,064 sequences were generated for CuSOD and MDH, respectively.

#### Ancestral sequence reconstruction

Maximum likelihood trees were generated from the training set reference MSAs using FastTree[22]. Ancestral sequence reconstructions were generated from the trees using the joint reconstruction function of the GRASP[23] command line tool. Metrics were calculated, and candidates were selected from the entire set of reconstructed sequences.

### Computational Metrics

#### AlphaFold2

AlphaFold2[24] was used to predict structures for test sequences and all generated sequences that passed the first filtering step. To create the features used by AlphaFold2, the complete database suit, including templates from the Protein data bank (pdb), was used. For each sequence, five predictions were made using the five models provided in AlphaFold2. The structure with the highest average pLDDT score was then selected as the representative structure for further analysis. AlphaFold2 predicted structures were used in the calculation of structure-based metrics, Rosetta-relax, ESM-IF, ProteinMPNN, and MIF-ST. The average pLDDT score, provided by AlphaFold2, across all residues was also used as a metric.

#### Phobius

The jphobius[9] (https://phobius.sbc.su.se/data.html) executable was used to predict the presence of signal peptides or transmembrane domains. The count of predicted transmembrane domains was recorded as an integer, and the presence or absence of a predicted signal peptide was recorded as a 1 or 0, respectively.

#### ESM-1v and CARP-640M

Scores calculated from the ESM-1v[25] and CARP-640M[26] models were the average of the log probabilities of the amino acid in each position. Without masking, this calculation can be done with a single forward pass over each sequence. With partial masking, it can be done in a number of passes equal to 1 / masked_fraction. At the most extreme, each position could be masked one at a time. We found that masking in six passes, with the masked positions at a regular interval that shifts on every pass, gave scores nearly equivalent to masking one position at a time. Furthermore, when no masking was applied, the scores were shifted towards zero but still strongly correlated with the masked scores. Therefore, we used ESM-1v and CARP-640M scores calculated in one pass without any masking.

#### ESM-MSA

Scores from the ESM-MSA-1b[17] model were calculated in a similar manner as for ESM-1v scores, the average log probability across the whole sequence. In the case of the ESM-MSA model, there were several hyperparameters to consider. Which sequences to include in the MSA, how many sequences to include in the MSA, how much to mask. We experimented with all of these hyperparameters. ESM-MSA scores are very sensitive to the composition of the MSA, but apparently less so to the size of the MSA.

For a masking strategy, we tried using unmasked, sparsely-masked, and fully-masked template sequences and found that sparse masking was the only strategy that gave satisfactory results. We settled on masking at regular intervals of 6 amino acids and therefore covering each entire sequence in six passes.

We first tried drawing reference sequences randomly from the training data. This gave high scores to sequences from areas of sequence space dense with training sequences and low scores to sequences from sparse areas of training sequence space, even at the same identity to the single closest training sequence. This effect could be mitigated by creating an alignment from the closest training sequences to each query.

We found that MSAs larger than about 32 gave ESM-MSA scores highly correlated to scores generated from larger MSAs.

Considering the above, our ESM-MSA metric was calculated by using phmmer[5] to find the 31 closest training sequences to each query, aligning the 32 sequences with MAFFT, and calculating the average log probabilities from six passes with a masking interval of six.

#### ProteinMPNN, ESM-IF, and MIF-ST

The proteinMPNN[27] and ESM-IF[28] scores are the average log-likelihood of the query residues, using the AlphaFold2 predicted structure. The ProteinMPNN score calculation is performed using the --score_only option of the protein_mpnn_run.py script from the ProteinMPNN repository (https://github.com/dauparas/ProteinMPNN). The ProteinMPNN score was multiplied by -1 so that higher is better, like with other metrics. The ESM-IF score is calculated using the esm.inverse_folding.util.score_sequence function from the ESM repository (https://github.com/facebookresearch/esm). The MIF-ST[29] score is calculated using the extract_mif.py script from the Protein Sequence Models repository (https://github.com/microsoft/protein-sequence-models).

#### Rosetta-relax

The relax program from Rosetta (v 2020.08.61146) [30] was used to relax the AlphaFold2 structures. The flags used for Rosetta relax were as follows:

```
-relax:constrain_relax_to_start_coords
-relax:ramp_constraints false
```

```
-ex1
-ex2
-use_input_sc
-no_optH false
-flip_HNQ
-nstruct 1
```

The total_score column was read from the output file, and Rosetta-relax score calculated according to the formula: -1 * (total_score / sequence_length). To normalize by sequence length and make it, so higher values for the metric correspond to proteins predicted to be more stable.

##### Distance to closest training sequence

The most similar training sequence was found using ggsearch36 from the FASTA package[31], the BLOSUM62 scoring matrix, a gap open penalty of 10, and a gap extend penalty of 2. The hamming distance was then calculated from the gapped alignment between the query and the top hit sequences. Identity was calculated as 1 - hamming_distance.

##### BLOSUM62 mutant position mean

The closest training sequence was found by ggsearch36 as described above. From the alignment to the closest training sequence, the mean BLOSUM62 score[32] across all mismatched positions was calculated, ignoring positions where either query or reference had a gap.

We considered that BLOSUM62 might not be the most appropriate matrix for our sequences, so we also calculated the alignments and scores using an alternative matrix, the PFASUM15 matrix [33], but the scores from the two matrices were highly correlated, thus we only report the BLOSUM62-based metric.

##### Longest repeat

Scores were calculated for the longest single amino acid repeat, the longest 2-mer, 3-mer, and 4-mer repeat in each sequence. The score was calculated as -1 * the number of repeat units. So the sequence AAAAAA would have a single amino acid repeat score of -6, a 2-mer score of -3, a 3-mer score of -2, and a 4-mer score of -1. The sequence LALALALA would have a 1-mer score of -1, a 2-mer score of -4, a 3-mer score of -1, and 4-mer score of -2.

#### Selection of sequences for in-vitro assays

##### Round 1

Sequences selected for experimental testing had between 70% and 80% identity to the closest training set sequence and diverse scores on the ESM-1v metric, representing the full range of scores found in generated sequences from that identity bin. Sequences were also filtered by manual inspection to remove those with large insertions or deletions compared to the closest reference sequences, or long repeats.

#### Round 2 pre-test

CuSOD sequences were manually selected based on the kingdom of origin, eukaryotic, viral, or bacterial, and the presence of Phobius predicted signal peptides. Sequences with predicted signal peptides were truncated at the predicted signal peptide cleavage site. Two bacterial FeSOD proteins, both lacking a predicted signal peptide, and the previously characterized[34] *E. coli* FeSOD (as a positive control) were also assayed.

#### Round 2

Sequences selected for experimental testing had between 80% and 90% identity to the closest training set sequence and diverse scores on the ESM-1v and ESM-MSA metrics, representing the full range of scores found in generated sequences from that identity bin. Sequences were also filtered by manual inspection to remove those with large insertions or deletions compared to the closest reference sequences or long repeats.

#### Round 3

Sequences were selected based on a series of filters. The first filter (which we call the "sequence filter" because it is based on sequence metrics) removed sequences having less than 50% or greater than 80% identity to the closest training sequence, having an ESM-1v score below the top 10 percentile threshold compared to the test sequences (-0.111 for CuSOD, -0.138 for MDH), sequences not starting with an M, having a predicted transmembrane domain, having a single amino acid repeat longer than 3 amino acids (for example, "AAAA"), or having an amino acid pair repeat longer than 4 amino acids (for example, "LALALA"), as repeats were more common in ESM-MSA generated sequences than in natural sequences (Supplementary Fig. 7). A total of 769,220 sequences were generated and scored in order to obtain at least 200 sequences passing the quality checks and ESM-1v threshold in the target identity band (50%-80%) for each model and enzyme. The numbers of sequences passing the filter were: 243 CuSOD-GAN, 371 CuSOD ESM-MSA, 441 MDH-GAN, 658 MDH-ESM-MSA. For each enzyme family, 200 ESM-MSA-generated sequences and 200 GAN-generated sequences were randomly selected from the sequences passing the first filter, and AlphaFold2 predicted their structure. ProteinMPNN scores were calculated for each of the structures, and the 40 sequences with the highest scores from each model/enzyme combination were kept. From the top 40 sequences, 18 were randomly selected for expression and functional characterization. For each passing sequence selected for functional characterization, a control sequence was selected from the list of sequences failing the sequence filter. Control sequences had an identity to the closest training sequence within 1% of the passing sequence and the longest single amino acid repeat of 6 or less (to facilitate gene synthesis, which struggles with very long repeats). A total of 144 sequences were synthesized and assayed in Round 3. 18 passing and 18 control sequences, each of GAN-generated CuSOD sequences, ESM-MSA generated CuSOD sequences, GAN-generated MDH sequences, and ESM-MSA generated MDH sequences.

For Round 3, we used an even more stringent cutoff, the top 10 percentile from the test set of natural sequences (the entire test set, not just the ones experimentally screened in Round 2).

### Experimental Assays

#### Bacterial strains, plasmids and growth conditions

*Escherichia coli* (*E.coli*) BL21(DE3) was used as the host strain for MDH and SOD expression host strain in this study. *E.coli* BL21(DE3) was normally grown on Luria-Bertaini (LB) medium at 37 °C supplemented with 100 μg/ml ampicillin (Cat#171254, Merck).
Amino acid sequences of MDH and SOD were converted into DNA sequences and codon-optimized for expression in *E.coli* on TWIST Bioscience (www.twistbioscience.com). A 30 bp sequence (TTTGTTTAACTTTAAGAAGGAGATATACAT) composed of RBS and spacer was added at the 5' terminus of all genes. All genes with 5' flank were cloned into the EcoRI and NotI sites of the pET-21(+) expression vector and sequence-verified by Twist Bioscience.
pET21b plasmid harboring MDH4 gene from a previous study [35] was used as a positive control for MDH enzymes. human SOD1[36] (hSOD, GenBank: NP_000445.1), *Potentilla atrosanguinea* (a plant) CuSOD[37] (paSOD, GenBank: AFN42318.1) and *E.coli* SOD [34] (E.SOD, GeneBank: NP_416173.1) were codon-optimized, synthesized in the same way as described above and used as positive controls for SOD enzymes. Blank plasmid pET21b was used as a negative control for both MDH and SOD enzymes.

#### Plasmid construction for truncated control sequences

All plasmids for truncated control sequences were prepared by subcloning the amplified sequences into the linearized pET21b plasmid (Cat#69741-3, Merck) by NdeI and NotI endogenous restriction enzymes with Gibson Assembly® Master Mix (Cat#E2611S, NEB). For hSOD, hSOD_Ntrun_F and hSOD_R primer pair was used to amplify the hSOD gene with 9 nucleotides (3 residues) truncated at the N terminus. hSOD_F and hSOD_Ctrun_R primer pair was used to amplify the human SOD1 gene with 12 nucleotides (4 residues) truncated at 3' terminus. hSOD_Ntrun_F and hSOD_Ctrun_R primer pair was used to amplify the human SOD1 gene with truncations at both ends. For paSOD, paSOD_Ntrun_F and paSOD_R primer pair was used to amplify the paSOD gene with 6 nucleotides (2 residues) truncated at N terminus. paSOD_F and paSOD_Ctrun_R primer pair was used to amplify the paSOD gene with 12 nucleotides (4 residues) truncated at 3' terminus. paSOD_Ntrun_F and paSOD_Ctrun_R primer pair was used to amplify the paSOD gene with truncations at both ends. E.coliSOD_F and E.coliSOD_Ctrun_R primer pair was used to amplify the E.SOD gene with truncations at 3' terminus. All primers used here were listed in Supplementary Table 7.

Insertions of genes were verified by colony PCR with T7_forward and T7term_reverse primer pairs (Supplementary Table 7). Plasmids from positive colonies were purified by GeneJET Plasmid Miniprep Kit (Cat#K0503, Thermo Scientific) and sequence-verified by Eurofins Genomics (www.eurofinsgenomics.eu).

#### Competent cell preparation and plasmid transformation

Competent cells of *E.coli* BL21(DE3) were prepared with the classic calcium chloride method [38]. Single colony of *E.coli* BL21(DE3) was inoculated into LB medium and grown overnight at 37 ℃ with 200 rpm shaking. 500 µl of the overnight grown culture was inoculated into 50 ml LB medium in a 250 ml shake flask. Cells were grown in 37 ℃ 200 rpm conditions until OD 600 reached 0.3 ~ 0.4. OD600 was monitored with a spectrophotometer (Genesys 20, Thermo Scientific). Cells were cooled down on ice and then spun down in a pre-cooled centrifuge with 4000 rpm for 10 min. The supernatant was discarded, and cell pellets were washed gently with 20 ml 0.1 M $CaCl_2$ once. After spinning down, cell pellets were suspended in 20 ml 0.1 M $CaCl_2$ and kept on ice for 30 min. After incubation, cells were spun down and suspended in 1 ml 0.1 M $CaCl_2$. 50 µl of suspension of competent cells was aliquoted into pre-chilled sterile 1.5 ml eppendorf tubes and continued to transformation or add glycerol to 15% final concentration for storage in - 80 ℃ until use.

20 µl MilliQ water was added into each well of lyophilized plasmids from the TWIST Bioscience well. 2 µl of plasmid solution was added into 50 µl competence cells and mixed well. The mixture was left on ice for 5 min and heat shocked in 42 ℃ water bath for 40 s. The tubes were left on ice for another 5 minutes, and then added 1 ml LB medium. The cells were shaken in 37 ℃ 200 rpm conditions for 1 h and then spread on LB agar plates supplemented with 100 µg/ml ampicillin. The plates were incubated at 37 ℃ overnight until single colonies grew out.

#### Protein expression and purification

Protein expression was achieved by diluting the overnight cultures 1:30 into 2.5 ml autoinduction Terrific Broth (TB) medium including trace elements (Cat#AIMTB0210, Formedium) and supplemented with 100 µg/ml ampicillin in 24-well format. All cells were cultivated in a 24-well plate in Eppendorf ThermoMixer® C. For MDH expression, cells were grown for 4 h at 37 °C, followed by overnight growth at 16 °C and 200 rpm shaking. For SOD expression, cells were grown for 4 h at 37 °C, followed by another 3 h at 25 °C and 200 rpm shaking.

Cells were collected by centrifugation at 5000 rpm for 10 min. Cell pellets were suspended in 200 µl Bugbuster (Cat#70584, Merck) supplemented with 1 µl 2000 U/ml DNaseI (Cat#79254, Qiagen) and incubated at 37 °C with 200 rpm shaking for 30 min. After incubation, 10 µl mixtures were aliquoted and kept in -20 °C as the sample of total protein (T) for gel electrophoresis. The mixture was centrifuged at maximum speed for 10 min, and pellets were discarded. 10 µl of the supernatant were aliquoted and kept at -20 °C as the sample of soluble protein (S) for gel electrophoresis. The supernatants were used for protein purification with the following procedures.

Talon resins (Cat#635653, Takara Bio) were washed with a binding buffer (50 mM $NaH_2PO_4$, 300 mM NaCl, 10 mM imidazole, pH 7.4) twice and then suspended in the same volume of a binding buffer as the resin bed amount. 50 µl Talon resin was loaded into each well of Nunc® 96 DeepWell™ filter plate (Cat#278012, Thermo Scientific). Each supernatant sample was added into the loaded column and then incubated at 4 °C for 30 min in a thermomixer.

The columns were then centrifuged at 200 x g for 30 s and flow waste was discarded. Resins were washed by 600 µl wash buffer for 3 times (50 mM $NaH_2PO_4$, 300 mM NaCl, 20 mM imidazole, pH 7.4) and centrifuged at 200 x g for 30 s each time. In the end, resins were incubated with 100 µl elution buffer at 4 °C for 30 min in a thermomixer and proteins were then eluted with 200 x g centrifugation for 1 min. Another 100 µl elution buffer was added to repeat the elution steps, and two portions of elution were mixed. The two eluate fractions were combined and transferred to a 96-well desalting plate (Cat#89807, Thermo Scientific), which was pre-equilibrated with sample buffer (50 mM $NaH_2PO_4$, 300 mM NaCl, pH 7.4). Protein samples were kept in -80 °C after adding 1x protein stabilizing cocktail (Cat#89806, Thermo Scientific). 10 µl of the proteins were aliquoted and kept at -20 °C as the sample of purified protein (P) for gel electrophoresis.

#### Gel electrophoresis

Total, soluble and purified proteins of each sample were mixed with 1x loading buffer (4x loading buffer recipe: 0.2 M Tris-HCl, 0.4 M DTT, 277 mM SDS, 6 mM Bromophenol blue, 4.3 M glycerol) and then heated in 85 °C for 5 min in a PCR cycler. Denatured proteins were analyzed by sodium dodecyl sulfate polyacrylamide gel electrophoresis (SDS–PAGE) with precast gels (Cat#WG1403A, Thermo Scientific), followed by coomassie staining with InstantBlue (Cat#ISB1L-53, Kem-en-tec). Spectra multicolor broad range protein ladder (Cat#26634, Thermo Scientific) was also loaded to analyze the protein sizes.

#### Enzymatic assay

To test for MDH activity, 2 µl of purified protein was added to a reaction mixture containing around 1.5 mM NADH (Cat#10128023001, Merck), 2.0 mM oxaloacetic acid (Cat#O4126, Sigma) and 20 mM HEPES buffer (pH 7.4). Assays were performed in triplicate and in 96-well format. All components were added by multichannel pipette to avoid the reaction time lag of each well. The final reaction volume was 100 µl, and the reaction was carried out at room temperature in a transparent 96-well microplate (Cat#0020821, Sarstedt) or 50 µl reaction volume in a transparent 384-well microplate (Nunc maxisorp, Cat#P6366, Merck). The activity was measured in triplicates by following NADH oxidation to NAD+, with an absorbance reading at 340 nm performed in kinetics mode for 15 min in a BMG Labtech SPECTROstar Nano spectrophotometer. Unspecific oxidation of NADH was monitored in no-substrate controls, and these values were subtracted from the other samples. Conversion from absorption values to NADH concentration was carried out using Beer-Lambert law $c=A/(d*\varepsilon)$, in which an extinction coefficient ε value is 6.22 $mM\cdot cm^{-1,}$ and path length for 100 µl in 96-well plate (d) is 0.29 cm. For those that did not show any catalytic activities, a 10-fold volume, which is 20 µl of purified proteins was used to perform the assay for a second time.
SOD activity was measured with SOD assay kit (Cat#19160, Sigma) in 96-well format, and all components were added by multichannel pipette to avoid the reaction time lag of each well. An aliquot (2 µl) of purified protein was added to each well containing 98 µl working solution. Assays of each sample were performed in triplicates and 1 “No XO” well. 10 µL of xanthine oxidase working solution was added into each well in the end, except for “No XO” wells. “No SOD” and “Blank” assays were also performed in triplicates. “No SOD” contained 10 µl dilution buffer, 80 µl working solution and 10 µL xanthine oxidase working solution, while “blank” contained 20 µl dilution buffer and 80 µl working solution. Plates were incubated in the plate reader, which was

pre-set at 37 °C. Absorbance at 450 nm was measured in kinetics mode for 30 min. For those that did not show any catalytic activities, a 10-fold volume, which is 20 μl of purified proteins, was used to perform the assay for a second time.
For the assay of truncated proteins, concentrations of desalted purified proteins were measured by Qubit Protein Assay kit (Cat#Q33211, Thermo Scientific) and 5 nM of all samples were used in the enzymatic assay.

#### Data analysis

For MDH, the absorbance value was plotted over time. The absorbance values of all samples at the endpoint of the assay were compared with negative control by T-test analysis. Samples were considered active if the end absorbance value was significantly lower than the negative control, i.e., $P$ value $\leq 0.05$.
For SOD, enzyme activity is measured as the percent inhibition of the rate of WST-1 formazan formation and calculated using the following equation with absorbance value at the time point of 20 min. The inhibition rate was compared with negative control by T-test, and those significantly higher than negative control were considered active with $P$ value $\leq 0.05$.

SOD activity (inhibition rate %) = [(A - B) – (C - D)]/ (A - B) x 100

A = Absorbance value of "No SOD" control
B = Absorbance value of Blank
C = Absorbance value of sample
D = Absorbance value of "No XO".

Assay data was analyzed with GraphPad Prism version 8.0.0 for Windows, GraphPad Software, San Diego, California USA, www.graphpad.com.

### Supplementary Figures

a

| | | Total | Active | Percent active |
|---|---|---|---|---|
| Kingdom | Eukaryote (CuSOD) | 4 | 3 | 75 |
| | Virus (CuSOD) | 1 | 1 | 100 |
| | Bacteria (CuSOD) | 9 | 4 | 44 |
| | Bacteria (FeSOD) | 2 | 2 | 100 |
| Signal peptide (CuSOD) | Clipped | 7 | 3 | 43 |
| | Never present | 7 | 4 | 57 |

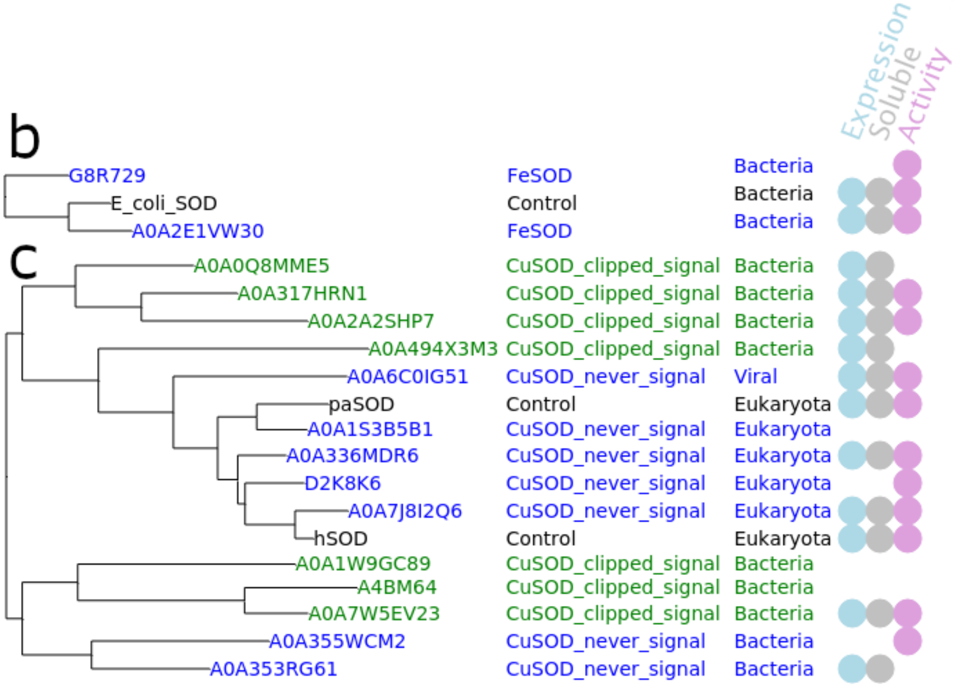


**Supplementary Fig 1. Round 2 pre-test assay results**. **a)** Table summarizing the results. **b)** Tree showing FeSOD results. **c)** Tree showing the CuSOD results. Controls (black), full-length sequences with no predicted signal peptide (blue), and sequences with N-terminal truncations to remove the predicted signal peptide (green). Circles indicate that the sequence was expressed, was soluble, or had activity.

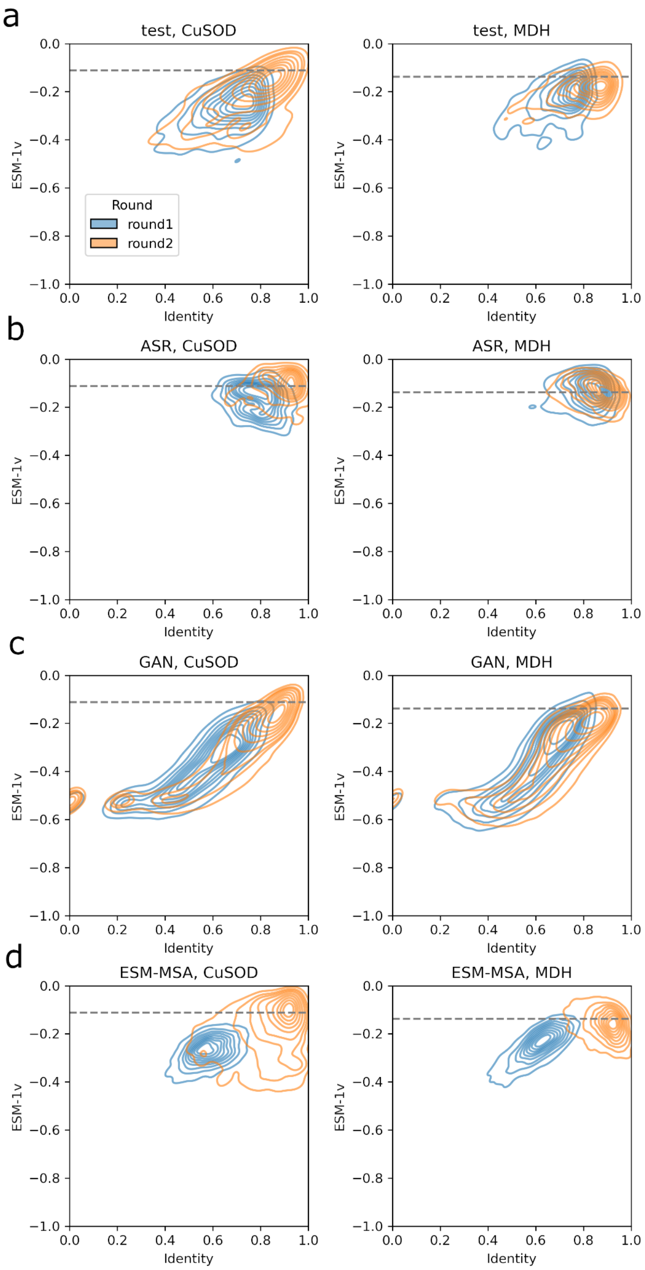


**Supplementary Fig 2. Density plots of Identity to most similar training sequence vs ESM-1v score for generated sequences from Rounds 1 and 2.** Dashed lines are the ESM-1v threshold used in the Round 3 filter. **a)** Natural sequences from the test set. **b)** Sequences generated by ancestral sequence reconstruction. **c)** Sequences generated by the GAN model. **d)** Sequences generated by the ESM-MSA model. The blue contours show the density of sequences generated for Round 1, the orange contours show the density of sequences generated for Rounds 2 and 3.

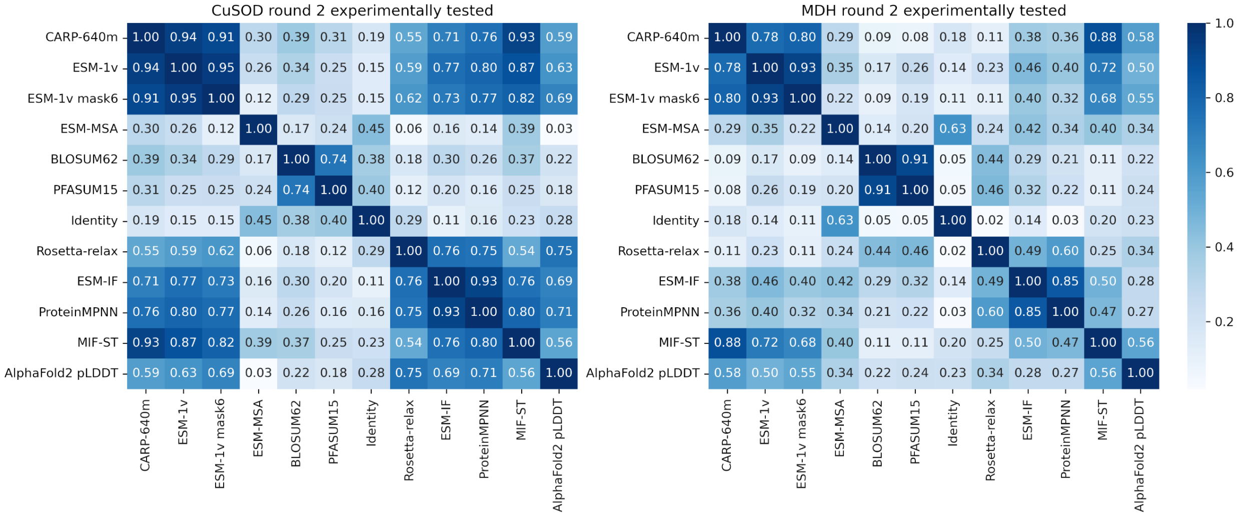


**Supplementary Fig 3. Spearman correlations between metrics for Round 2**. Experimentally tested CuSOD (left) and MDH (right) sequences.

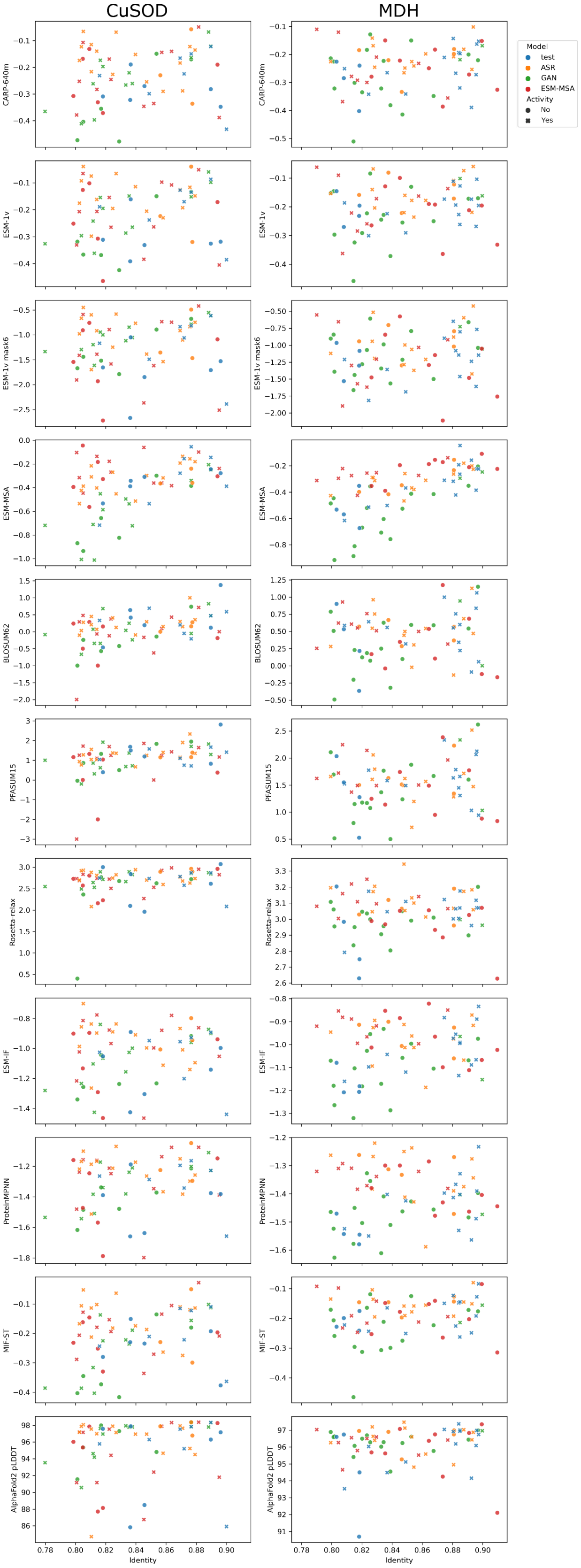


**Supplementary Fig 4. Plots of identity vs Round 2 metrics**

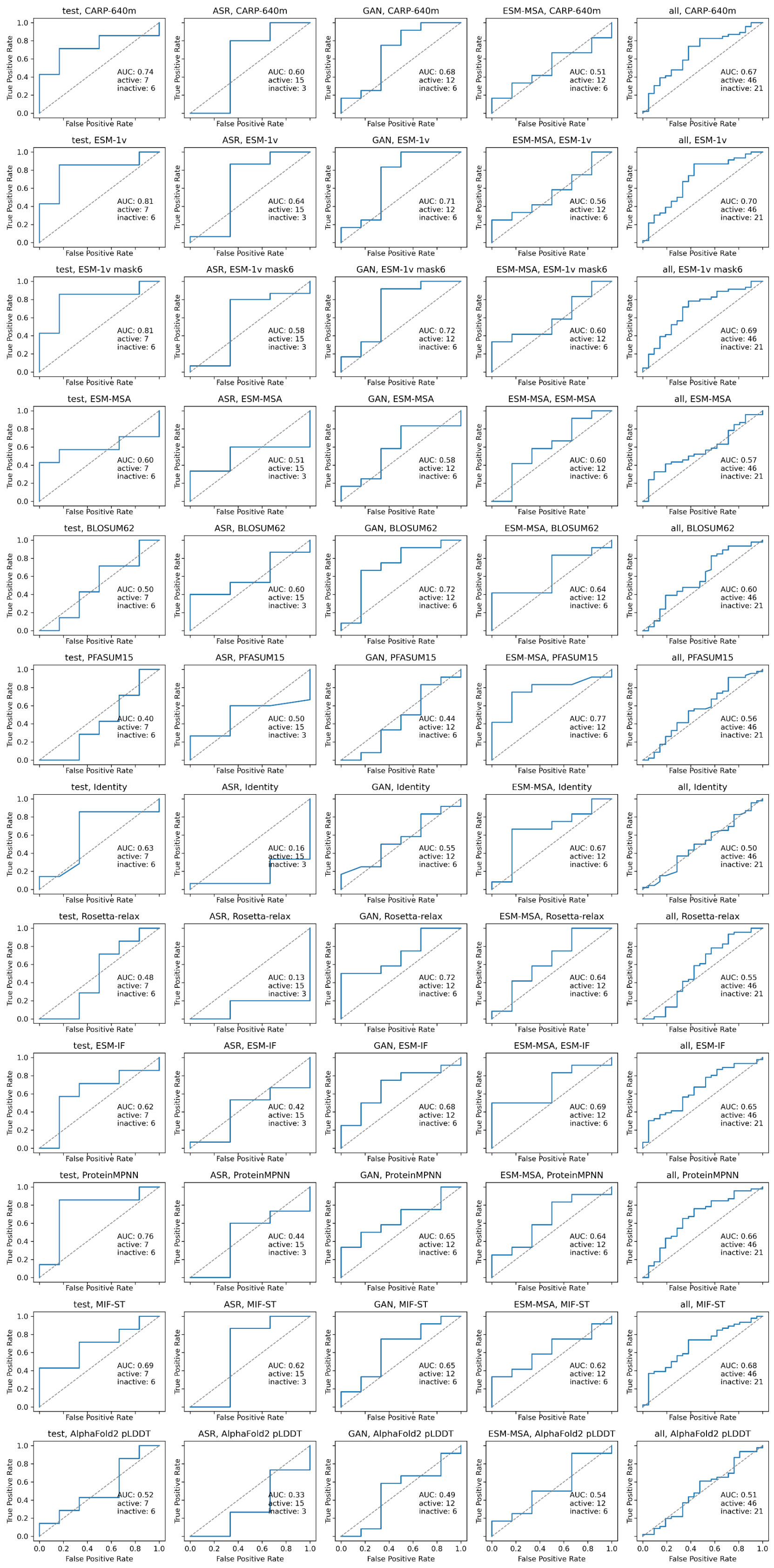


**Supplementary Fig 5. Activity vs metric ROC curves for Round 2 CuSOD enzymes**

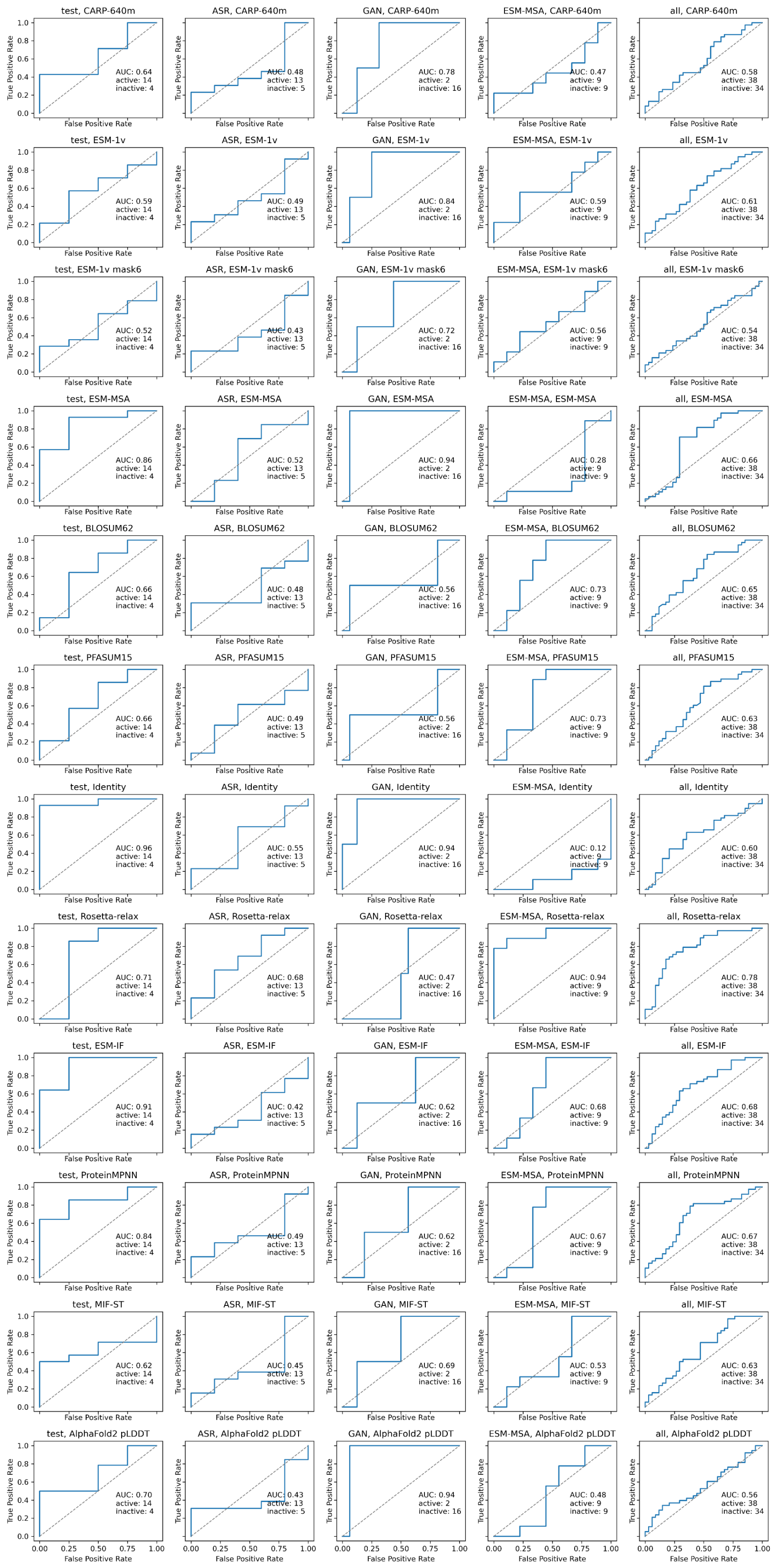


**Supplementary Fig 6. Activity vs metric ROC curves for Round 2 MDH enzymes**

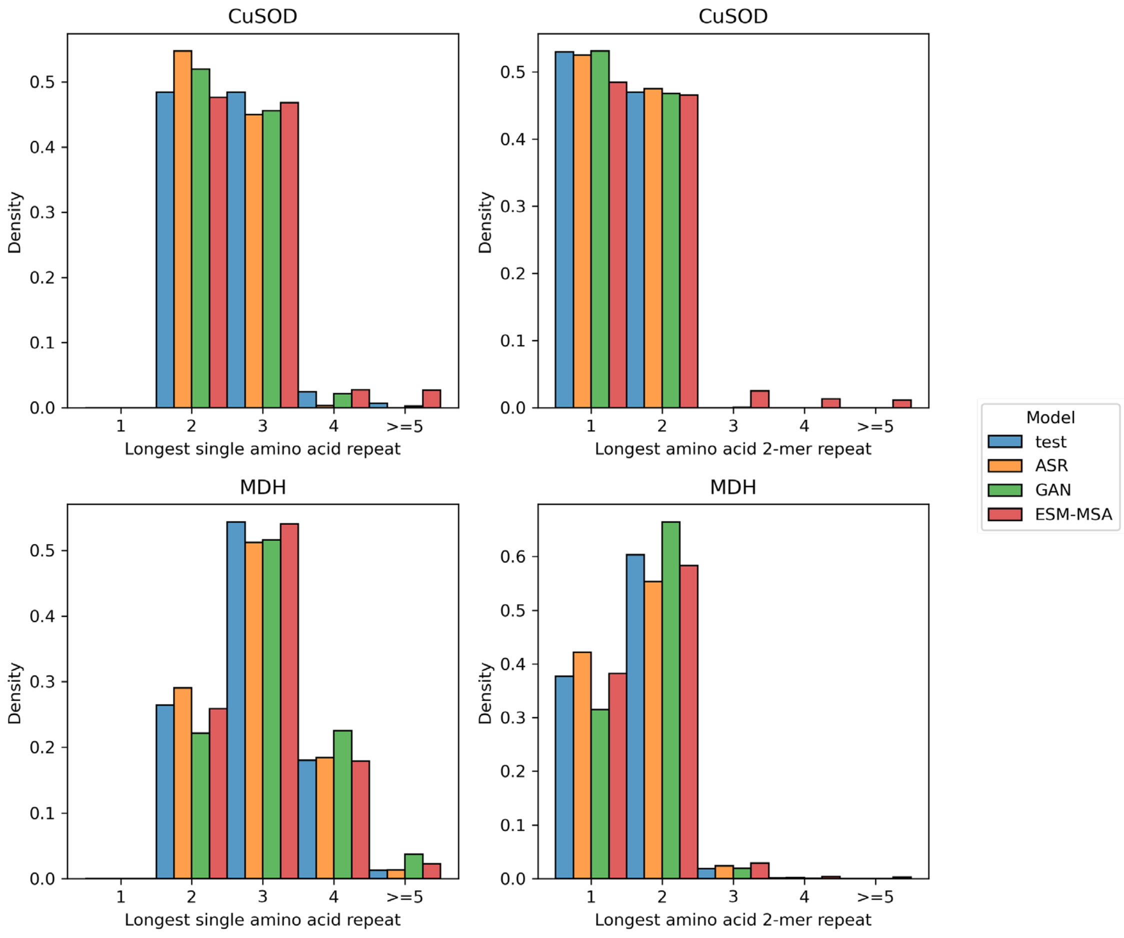


**Supplementary Fig 7. Longest repeat sizes per sequence in sequences generated for Rounds 2 and 3.**

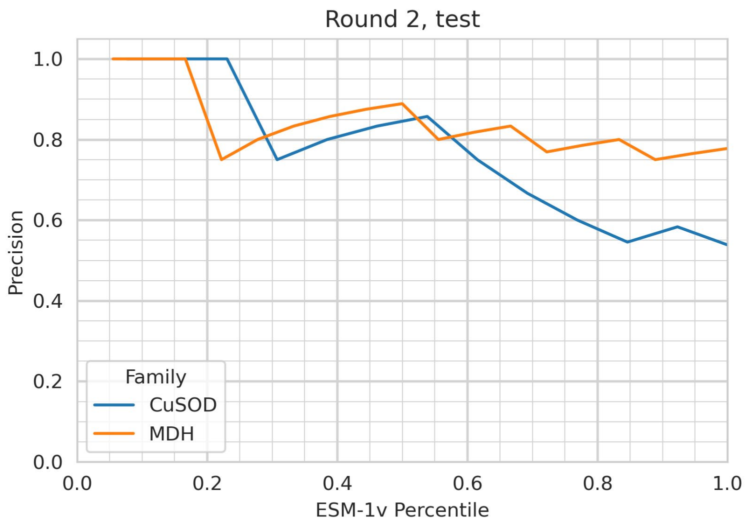


**Supplementary Fig 8. Precision vs ESM-1v percentile cutoff curves for activity in Round 2 test (natural) sequences.**

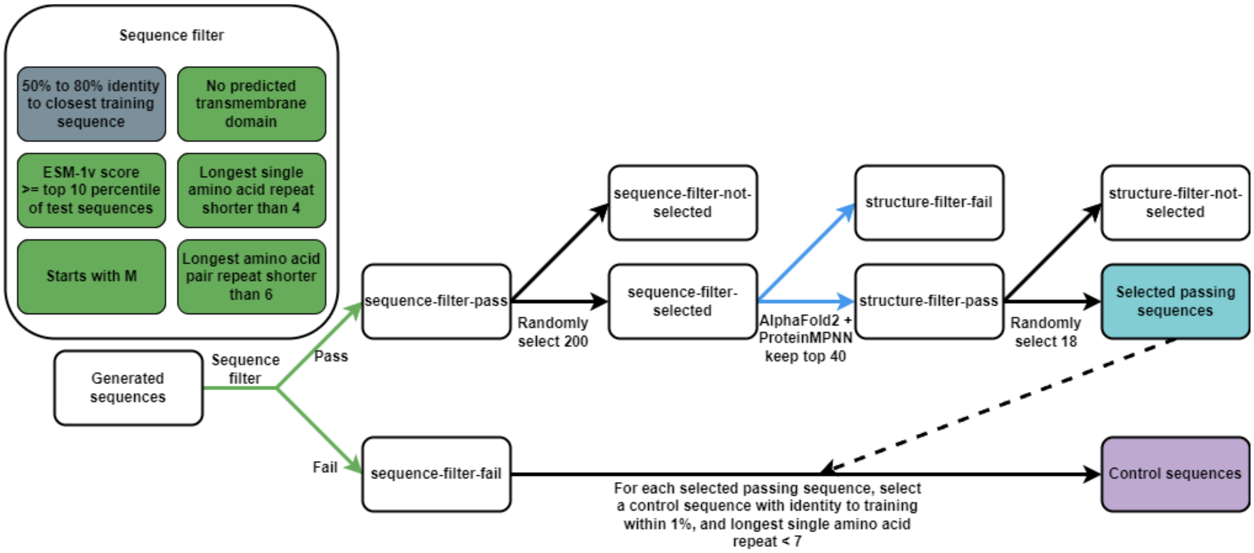


**Supplementary Fig 9. Expanded schematic for Round 3 selection process.**

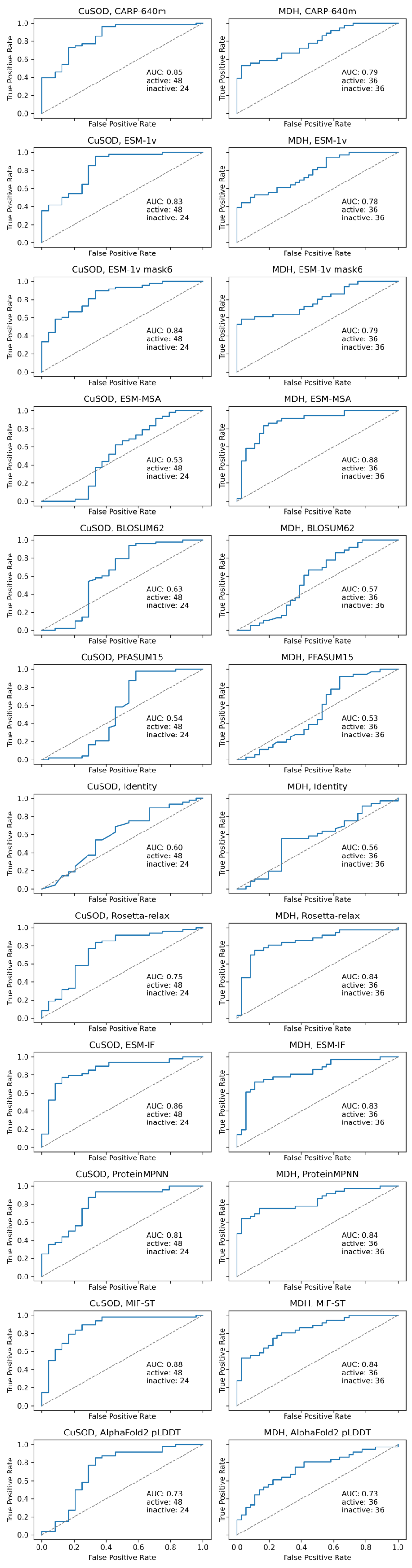


**Supplementary Fig 10. Activity vs metric ROC curves for Round 3.** CuSOD (left) and MDH (right) enzymes.

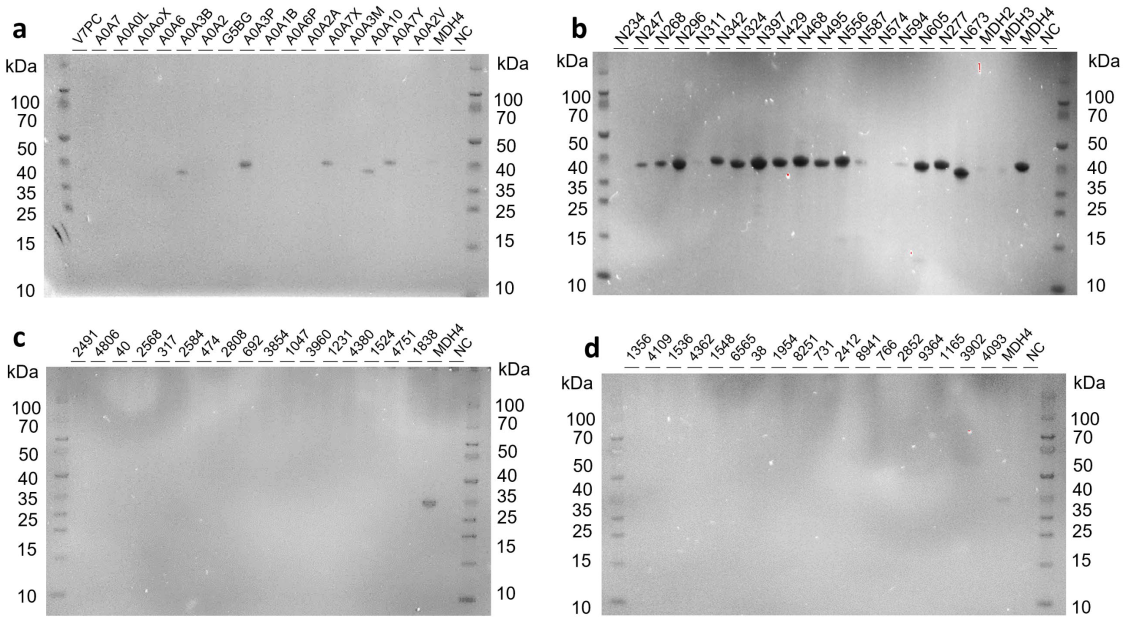

**Supplementary Fig 11. SDS gel of purified MDH enzymes in Round 1** from a) test group, b) ASR group, c) ESMMSA group and d) GAN group.

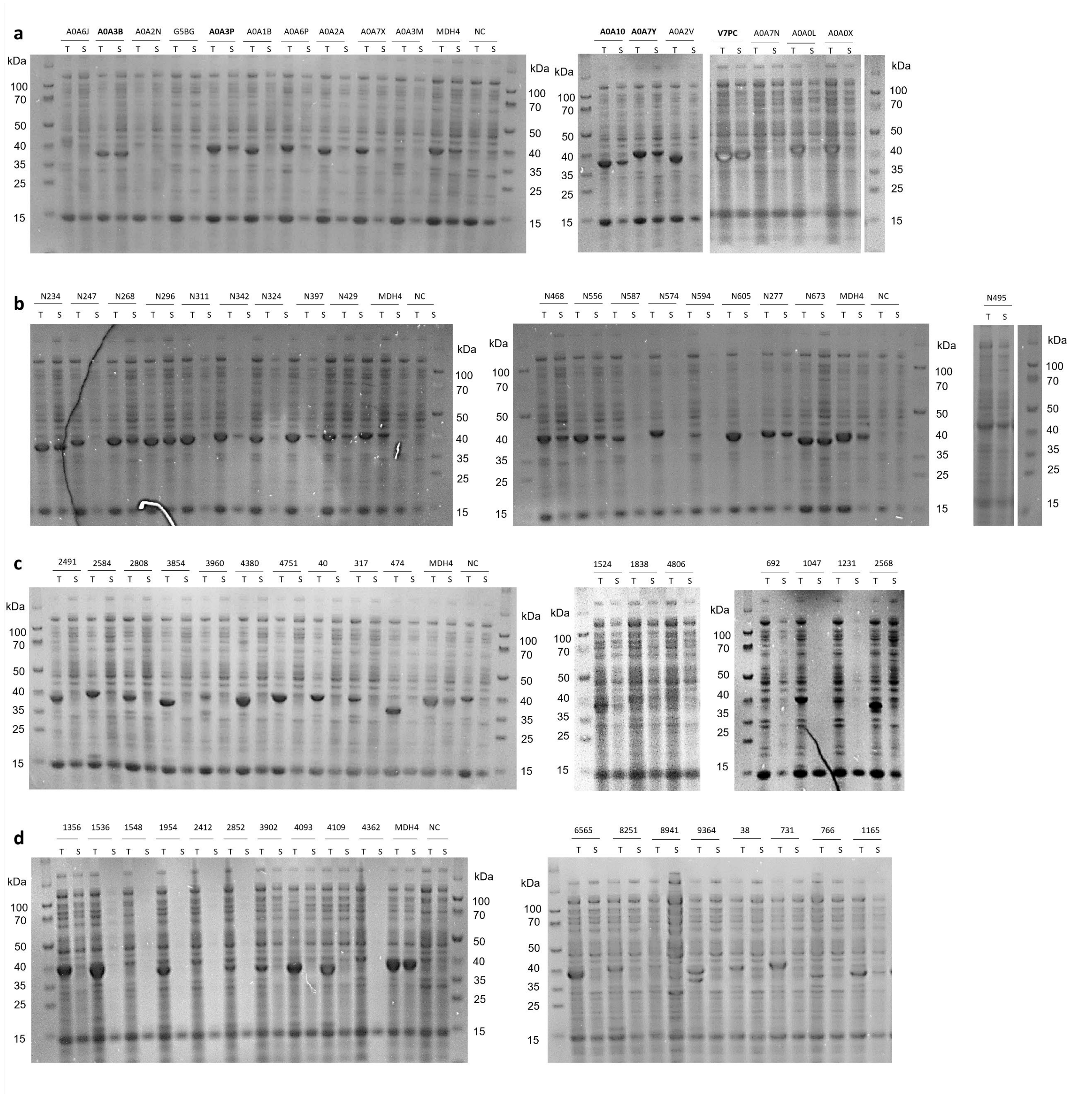


**Supplementary Fig 12. SDS gel of total (T) and soluble (S) MDH protein samples in Round 1** from a) test group, b) ASR group, c) ESMMSA group and d) GAN group.

a

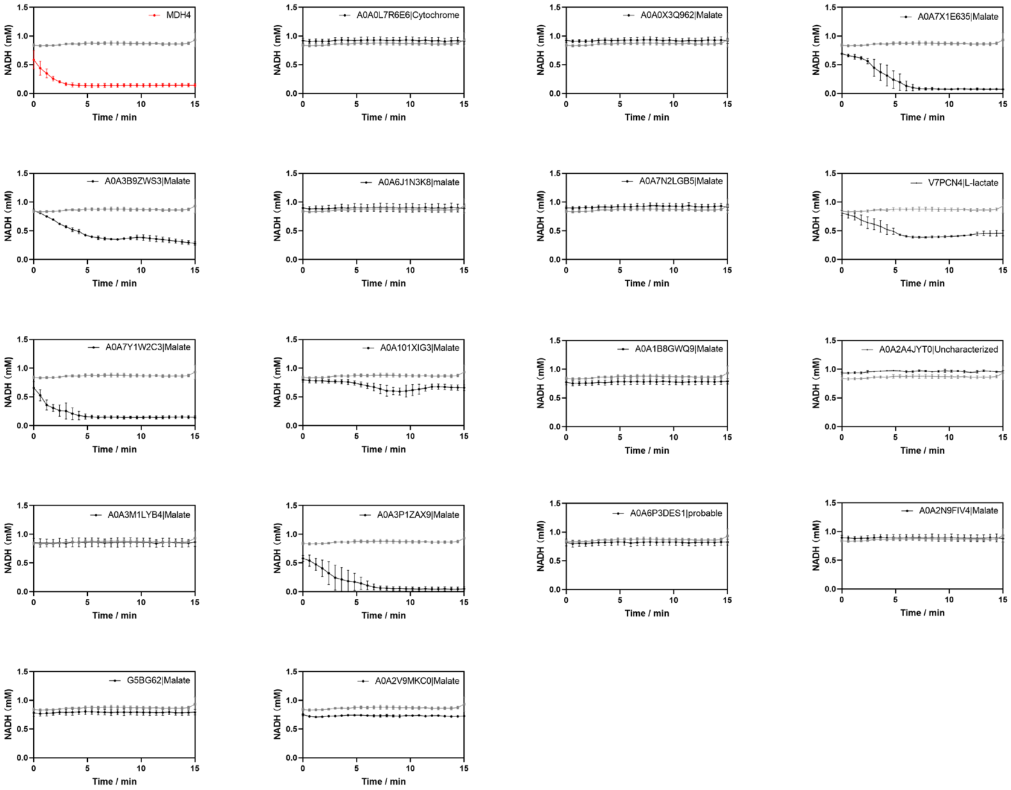
MDH4
A0A0L7R6E6|Cytochrome
A0A0X3Q962|Malate
A0A7X1E635|Malate
A0A3B9ZWS3|Malate
A0A6J1N3K8|malate
A0A7N2LGB5|Malate
V7PCN4|L-lactate
A0A7Y1W2C3|Malate
A0A101XIG3|Malate
A0A1B8GWQ9|Malate
A0A2A4JYT0|Uncharacterized
A0A3M1LYB4|Malate
A0A3P1ZAX9|Malate
A0A6P3DES1|probable
A0A2N9FIV4|Malate
G5BG62|Malate
A0A2V9MKC0|Malate
NADH (mM)
Time / min

b

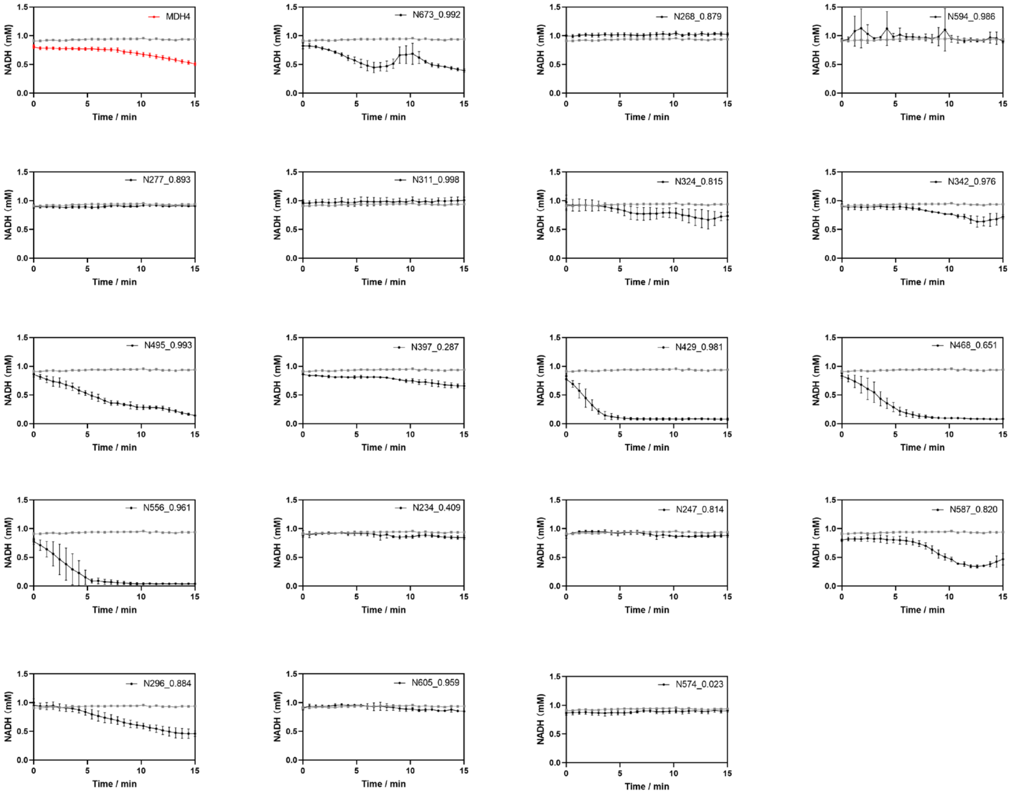
MDH4
N673_0.992
N268_0.879
N594_0.986
N277_0.893
N311_0.998
N324_0.815
N342_0.976
N495_0.993
N397_0.287
N429_0.981
N468_0.651
N556_0.961
N234_0.409
N247_0.814
N587_0.820
N296_0.884
N605_0.959
N574_0.023
NADH (mM)
Time / min

**c**

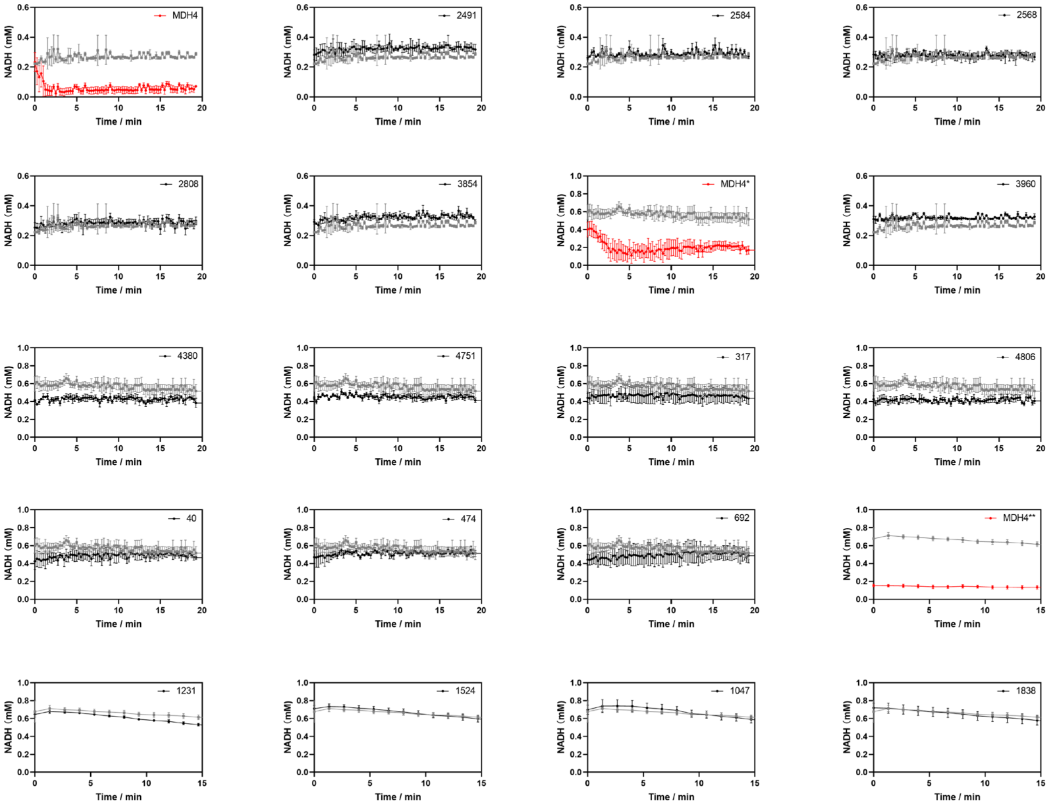
NADH (mM)
Time / min
MDH4
2491
2584
2568
2808
3854
MDH4*
3960
4380
4751
317
4806
40
474
692
MDH4**
1231
1524
1047
1838

**d**

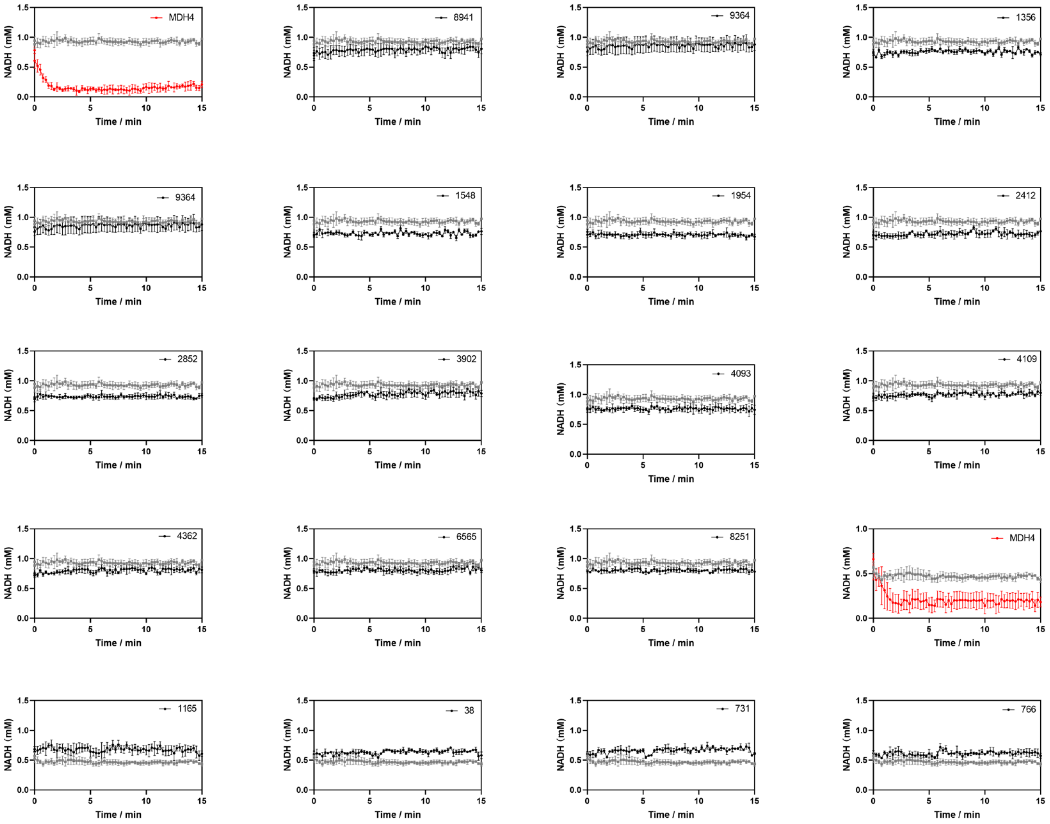
NADH (mM)
Time / min
MDH4
8941
9364
1356
9364
1548
1954
2412
2852
3902
4093
4109
4362
6565
8251
MDH4
1165
38
731
766

**Supplementary Fig 13. MDH enzyme activities in Round 1** from a) test group, b) ASR group, c) ESMMSA group and d). GAN group measured in *vitro* by monitoring NADH consumption. An *E. coli* strain harboring pET-21b was used as a negative control. Wild-type malate dehydrogenases (MDH 4), likewise produced in *E. coli*. serves as positive controls. A time-dependent decrease in NADH can be readily observed for moderately active enzymes. Due to the limitation of 96 well format, assays were performed in several batches and both positive and negative controls were included in each batch of assay. For highly active enzymes all the NADH was consumed before measurements could take place, resulting in a horizontal line at the bottom of the plot. Negative controls are plotted in grey, positive controls in red, and generated sequences in black.

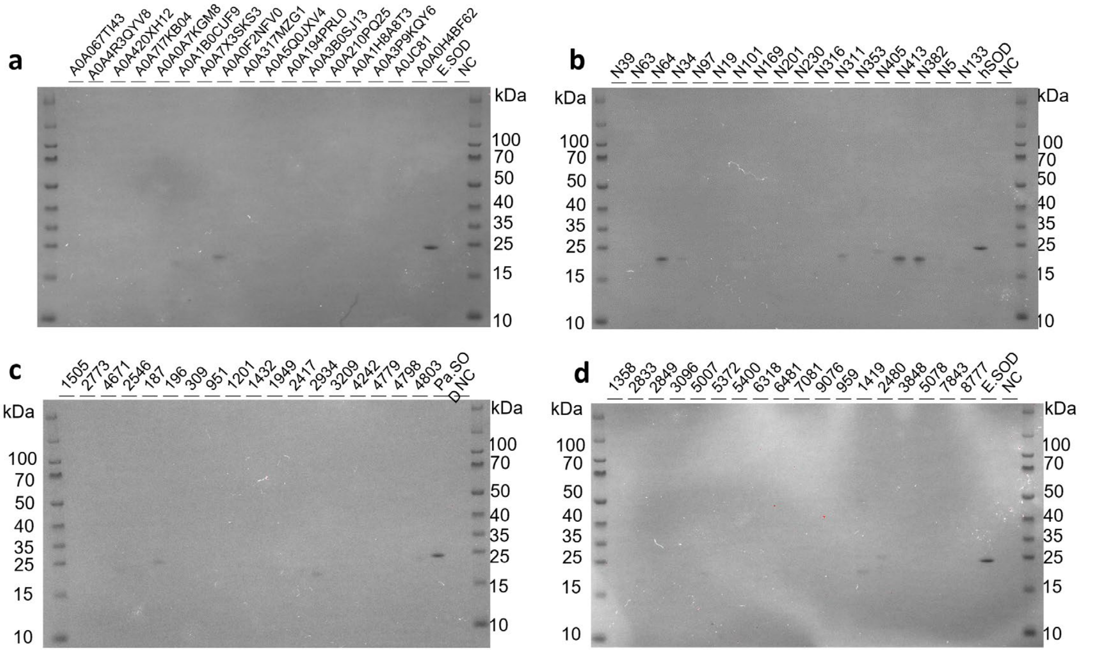


**Supplementary Fig 14**. **SDS gel of purified SOD enzymes in Round 1** from a) test group, b) ASR group, c) ESMMSA group and d). GAN group.

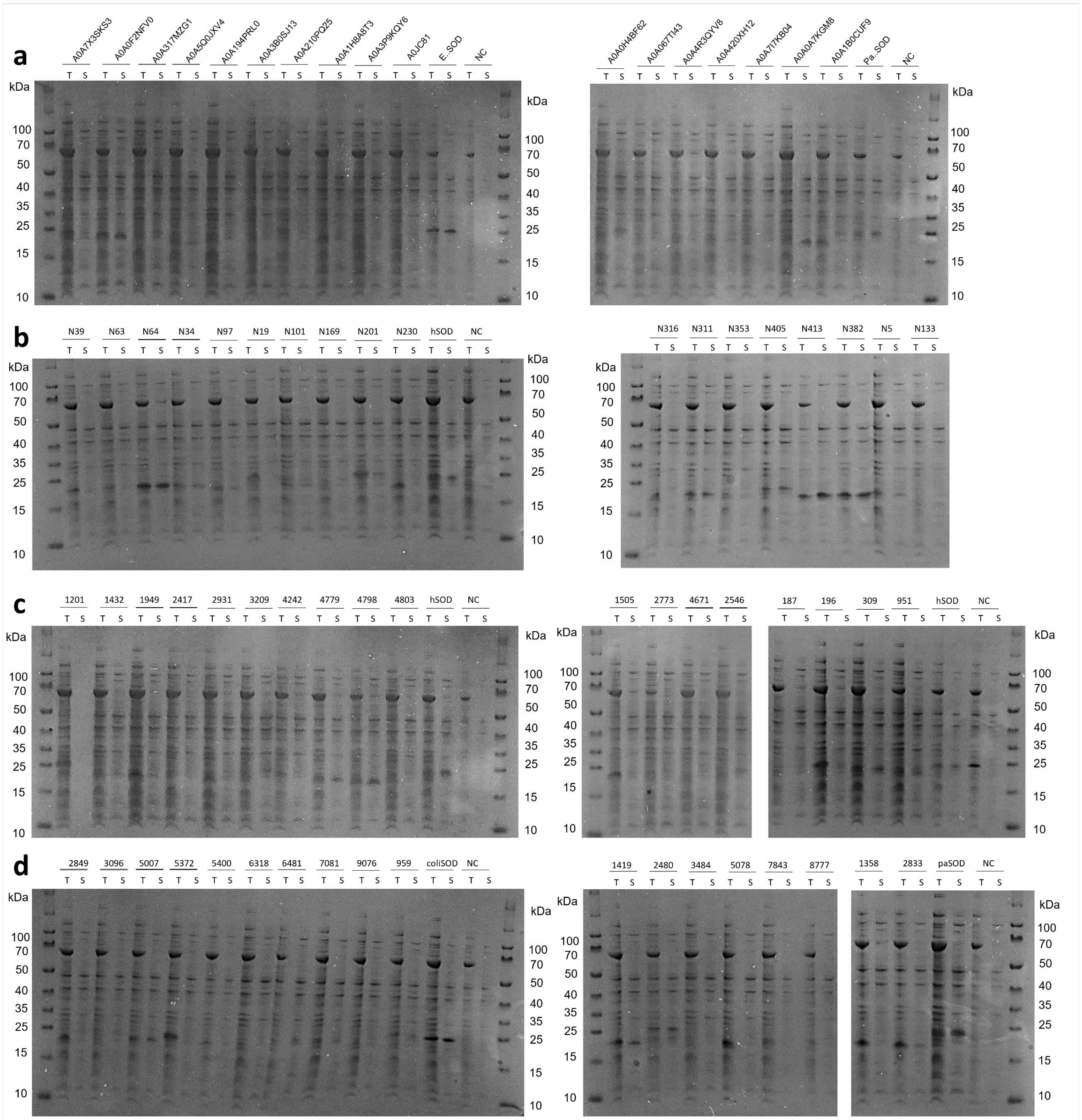


**Supplementary Fig 15. SDS gels of total (T) and soluble (S) SOD protein samples in Round 1** from a) test group, b) ASR group, c) ESMMSA group and d) GAN group.

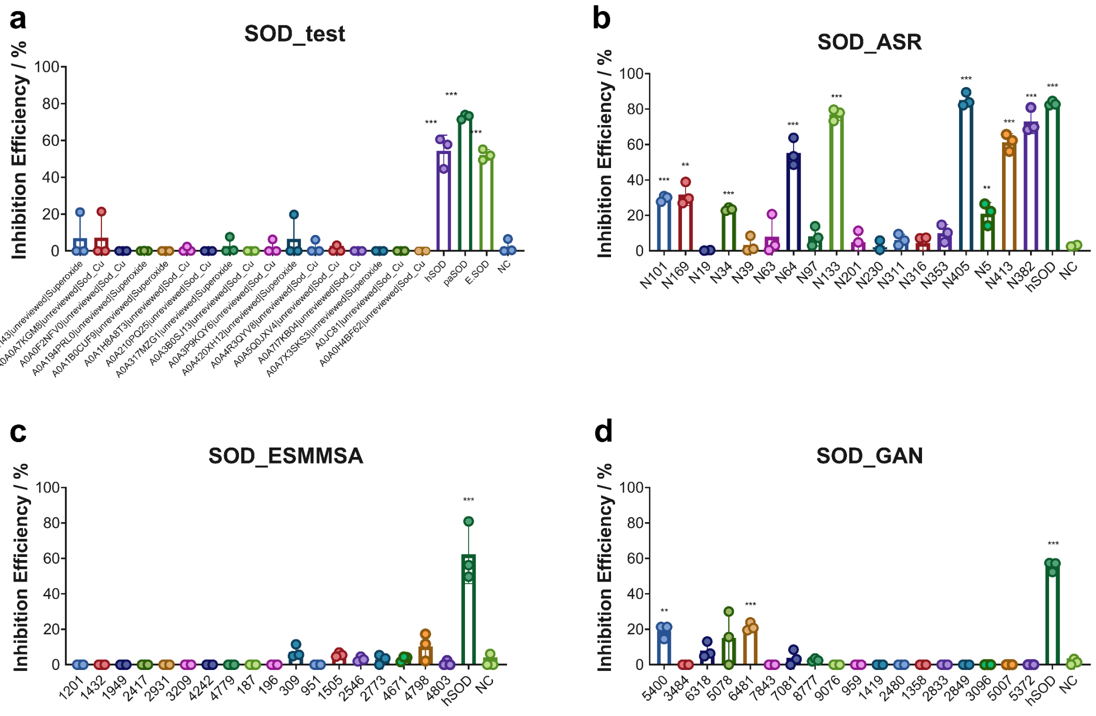


**Supplementary Fig 16. SOD enzyme activities in Round 1** from a) test group, b) ASR group, c) ESMMSA group and d) GAN group, which measures *in vitro* activity by the inhibition efficiency on xanthine oxidase. * marks samples with P-value less than 0.05 from the t-test compared to the negative control samples, ** for P < 0.01 and *** for P < 0.001.

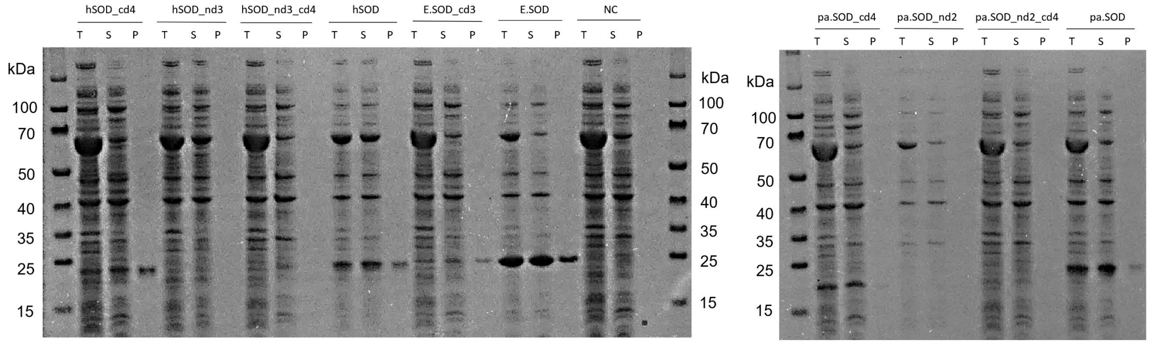


**Supplementary Fig 17. SDS gel of total (T), soluble (S) and purified (P) SOD protein in truncation experiments**.

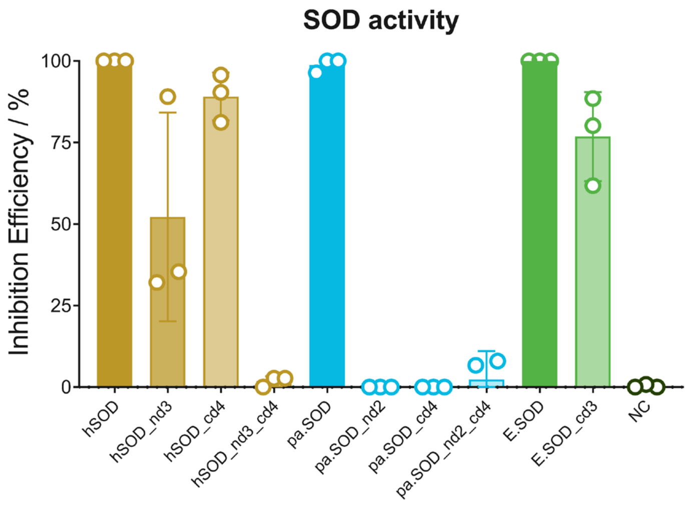


**Supplementary Fig 18. Enzymatic activity of truncated hSOD and paSOD** with truncations at N terminus (nd), C terminus (cd) and both ends, **E.SOD** with 3 amino acid truncation at C terminus, in comparison with intact hSOD, paSOD, E.SOD and negative control (NC).

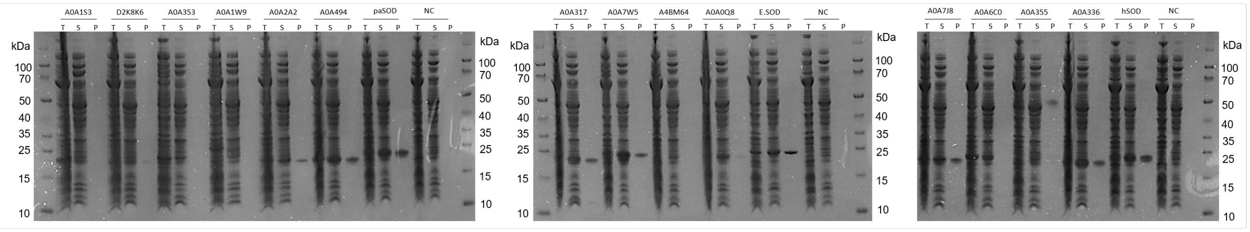


**Supplementary Fig 19. SDS gel of total (T), soluble (S) and purified (P) SOD protein samples in pre-test group.**

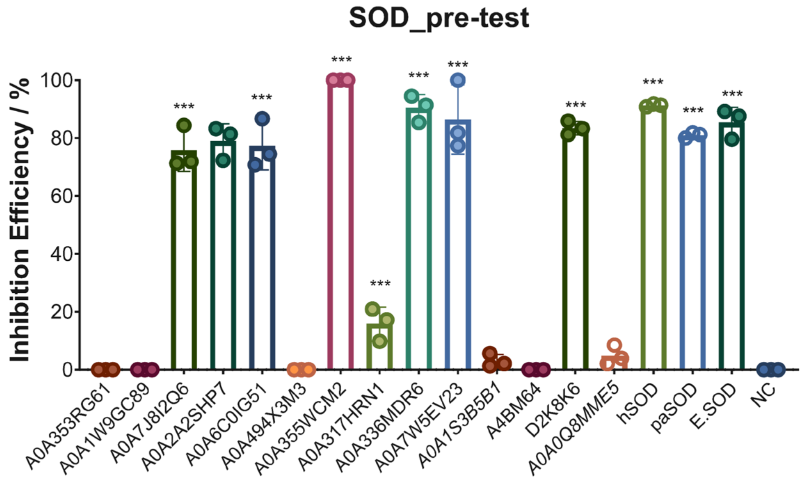


**Supplementary Fig 20. SOD enzyme activities in pre-test group.** SOD activity was measured *in vitro* indirectly by the inhibition efficiency on xanthine oxidase. * marks samples with P-value less than 0.05 from the t-test compared to the negative control samples, ** for P < 0.01 and *** for P < 0.001.

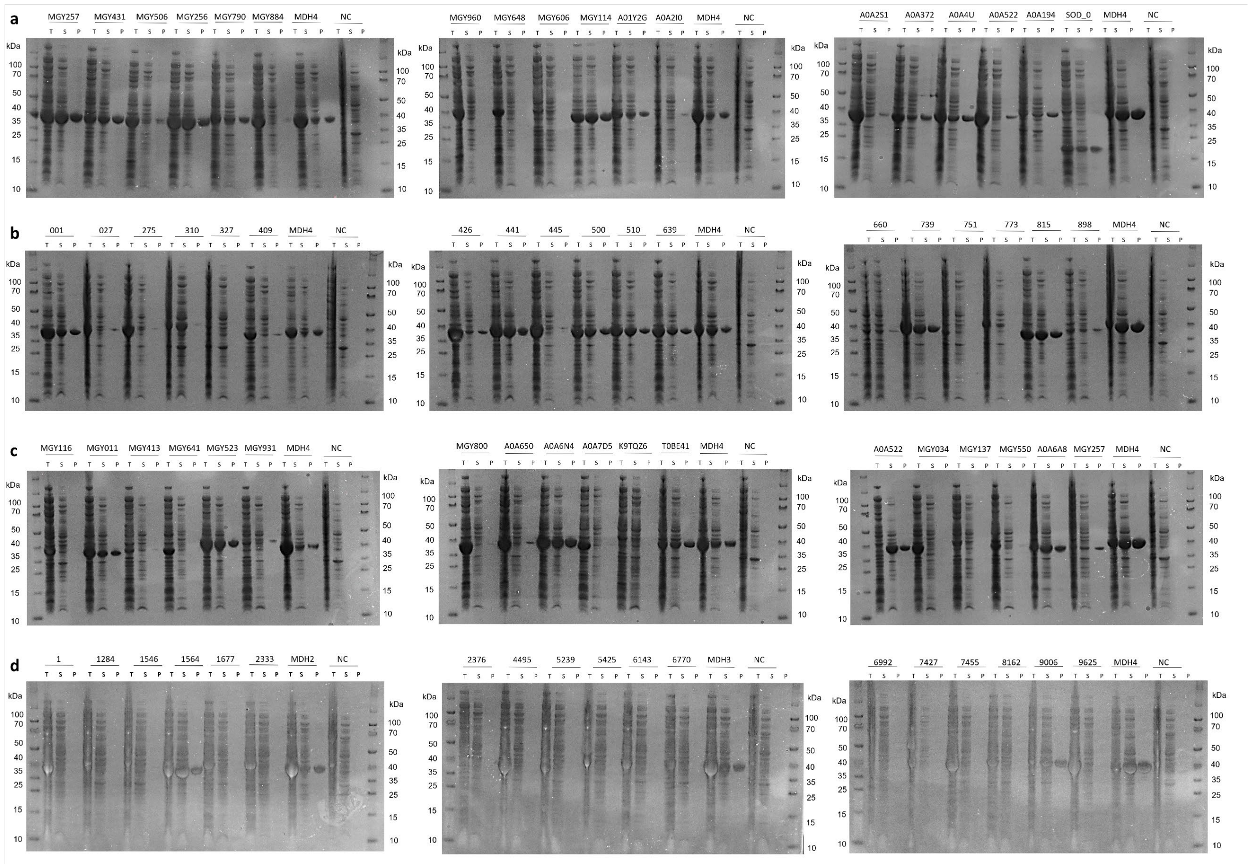

**Supplementary Fig 21. SDS gel of total (T), soluble (S) and purified (P) MDH protein samples in Round 2** from a) test group, b) ASR group, c) ESMMSA group and d) GAN group.

**a**

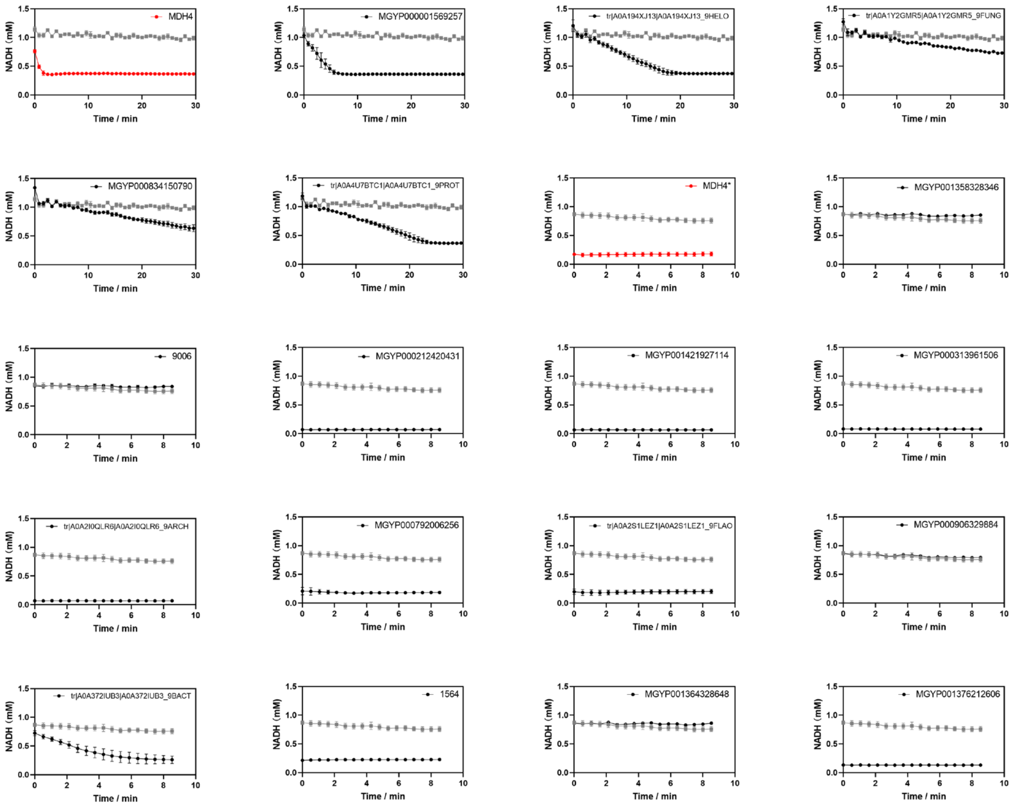

NADH (mM)
Time / min
MDH4
MGYP000001569257
tr|A0A194XJ13|A0A194XJ13_9HELO
tr|A0A1Y2GMR5|A0A1Y2GMR5_9FUNG
MGYP000834150790
tr|A0A4U7BTC1|A0A4U7BTC1_9PROT
MDH4*
MGYP001358328346
9006
MGYP000212420431
MGYP001421927114
MGYP000313961506
tr|A0A2I0QLR6|A0A2I0QLR6_9ARCH
MGYP000792006256
tr|A0A2S1LEZ1|A0A2S1LEZ1_9FLAO
MGYP000906329884
tr|A0A372IUB3|A0A372IUB3_9BACT
1564
MGYP001364328648
MGYP001376212606


**b**

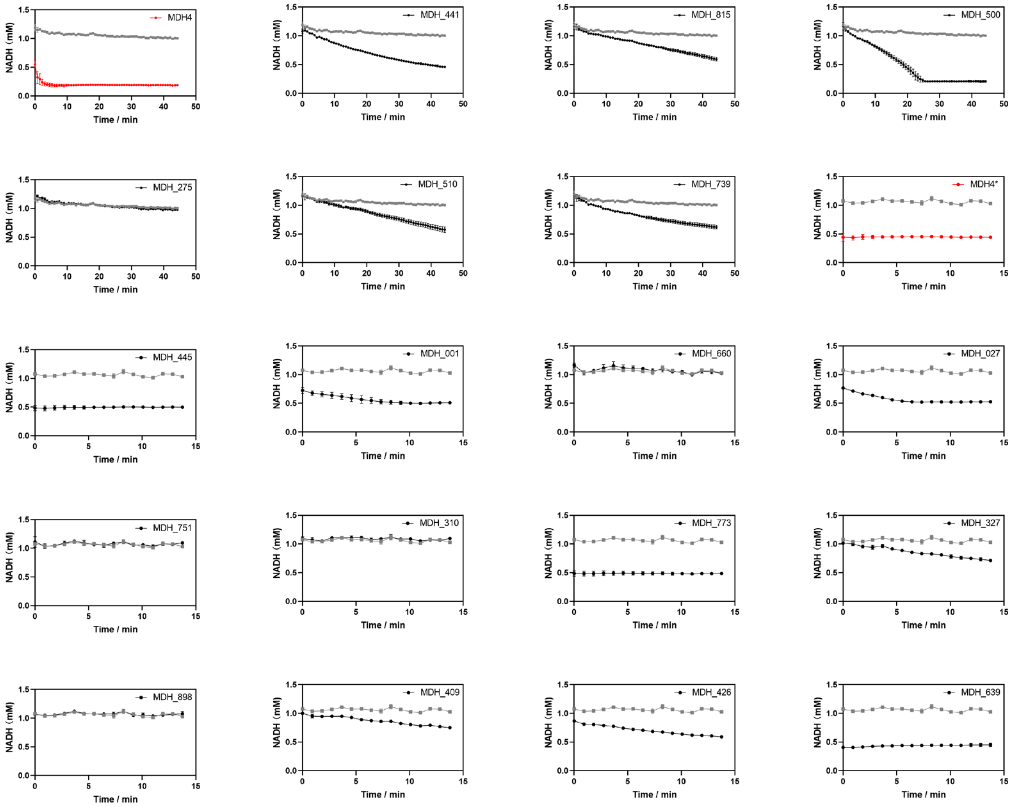

NADH (mM)
Time / min
MDH4
MDH_441
MDH_815
MDH_500
MDH_275
MDH_510
MDH_739
MDH4*
MDH_445
MDH_001
MDH_660
MDH_027
MDH_751
MDH_310
MDH_773
MDH_327
MDH_898
MDH_409
MDH_426
MDH_639

**c**

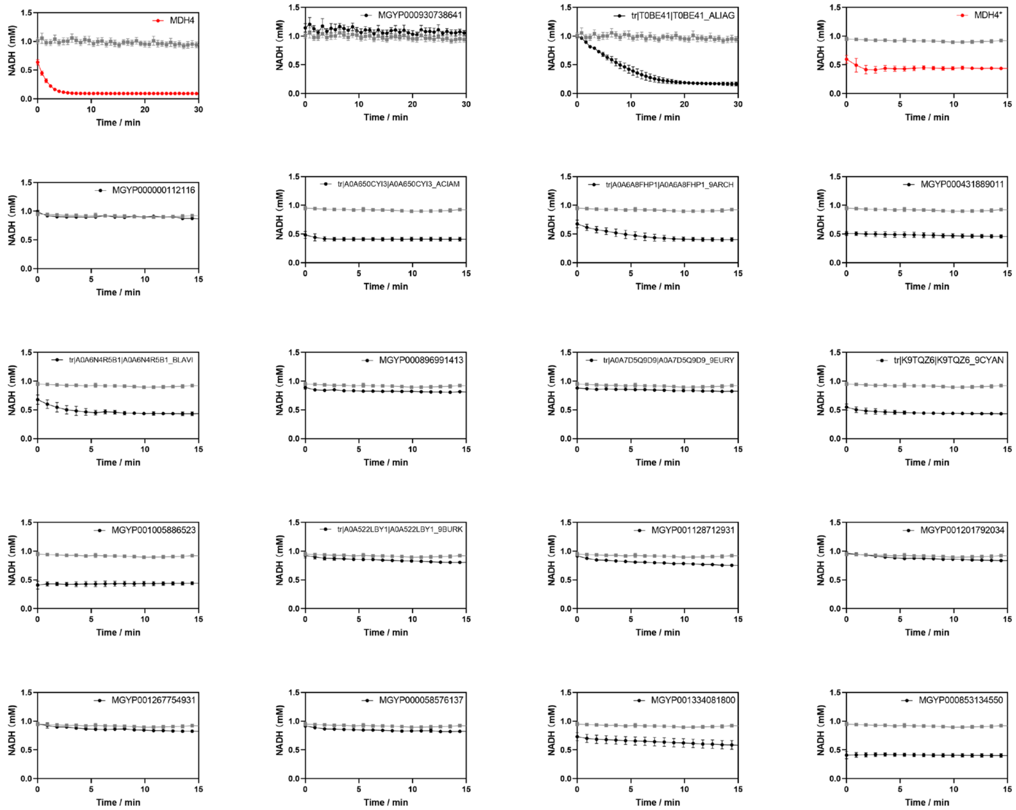

MDH4
MGYP000930738641
tr|T0BE41|T0BE41_ALIAG
MDH4*
MGYP000000112116
tr|A0A650CYI3|A0A650CYI3_ACIAM
tr|A0A6A8FHP1|A0A6A8FHP1_9ARCH
MGYP000431889011
tr|A0A6N4R5B1|A0A6N4R5B1_BLAVI
MGYP000896991413
tr|A0A7D5Q9D9|A0A7D5Q9D9_9EURY
tr|K9TQZ6|K9TQZ6_9CYAN
MGYP001005886523
tr|A0A522LBY1|A0A522LBY1_9BURK
MGYP001128712931
MGYP001201792034
MGYP001267754931
MGYP000058576137
MGYP001334081800
MGYP000853134550
NADH (mM)
Time / min


**d**

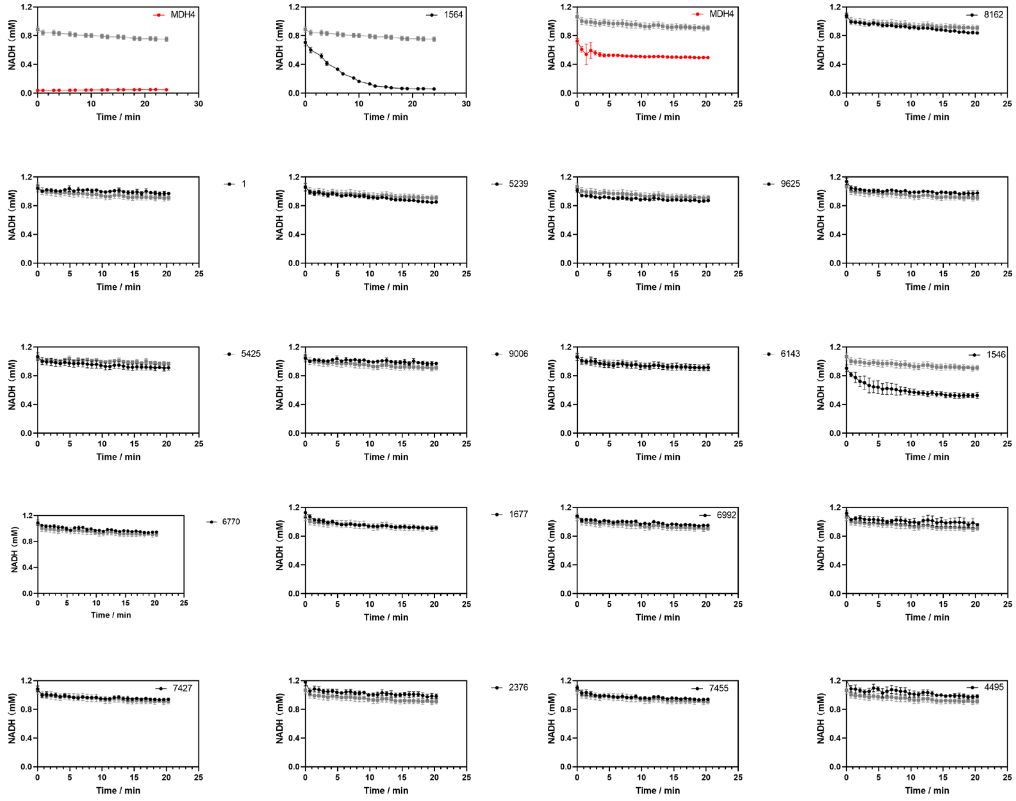

MDH4
1564
MDH4
8162
1
5239
9625
5425
9006
6143
1546
6770
1677
6992
7427
2376
7455
4495
NADH (mM)
Time / min

**Supplementary Fig 22. MDH enzyme activities in Round 2** from a) test group, b) ASR group, c) ESMMSA group and d) GAN group in Round 2 measured in *vitro* by monitoring NADH consumption. An *E. coli* strain harboring pET-21b was used as a negative control. Wild-type malate dehydrogenases (MDH 4), likewise produced in *E. coli*. serves as positive controls. A time-dependent decrease in NADH can be readily observed for moderately active enzymes. Due to the limitation of 96 well format, assays were performed in several batches and both positive and negative controls were included in each batch of assay. For highly active enzymes all the NADH was consumed before measurements could take place, resulting in a horizontal line at the bottom of the plot. Negative controls are plotted in grey, positive controls in red, and generated sequences in black.

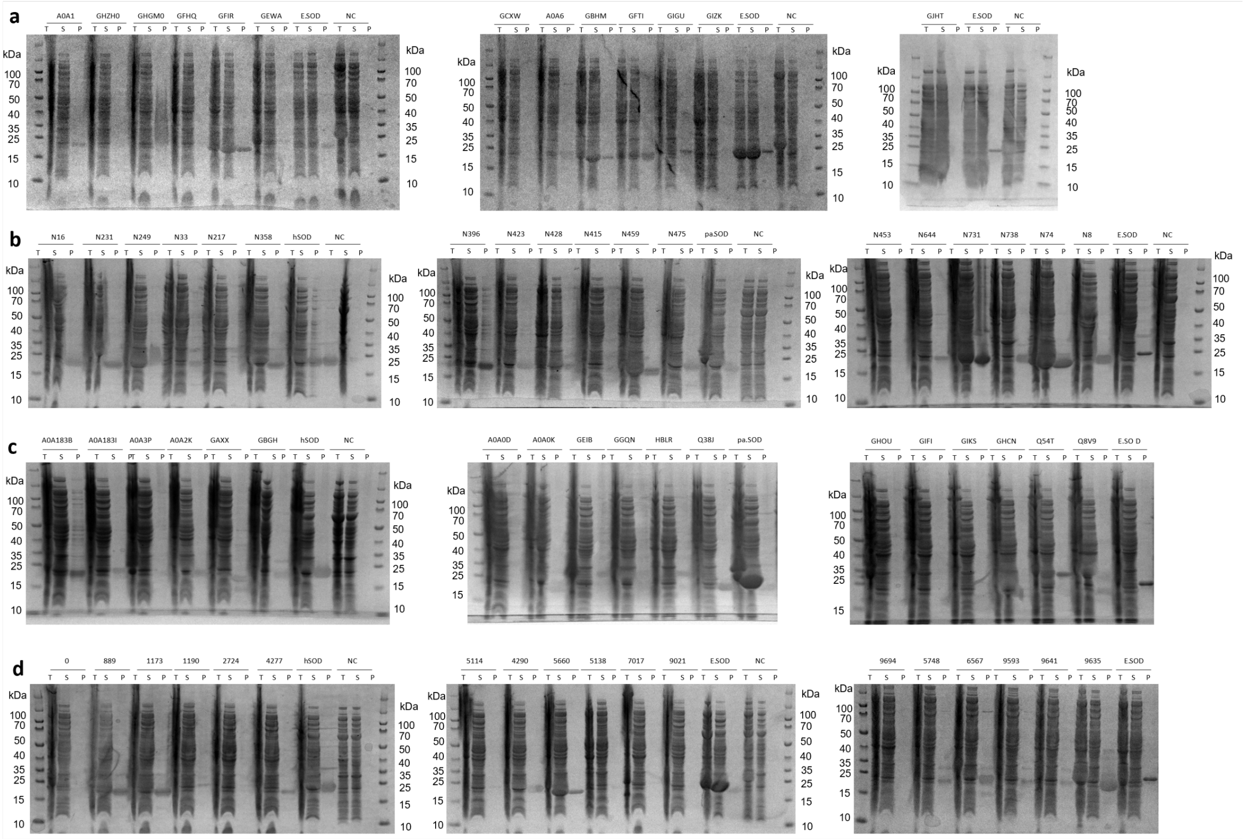

**Supplementary Fig 23. SDS gel of total (T), soluble (S) and purified (P) SOD protein samples in Round 2** from a) test group, b) ASR group, c) ESMMSA group and d) GAN group.

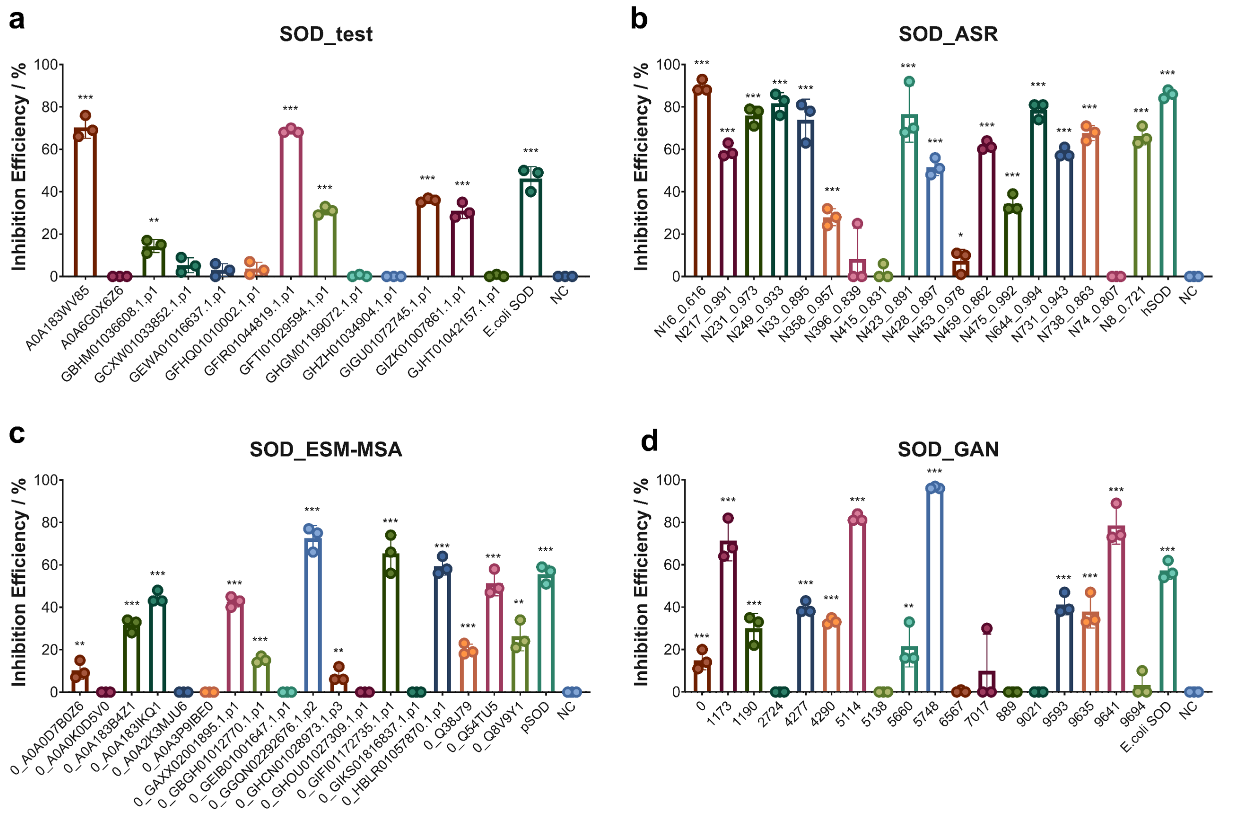


**Supplementary Fig 24. SOD enzyme activities in Round 2** from a) test group, b) ASR group, c) ESMMSA group and d) GAN group, which were measured *in vitro* activity by the inhibition efficiency on xanthine oxidase. * marks samples with P-value less than 0.05 from the t-test compared to the negative control samples, ** for $P < 0.01$ and *** for $P < 0.001$.

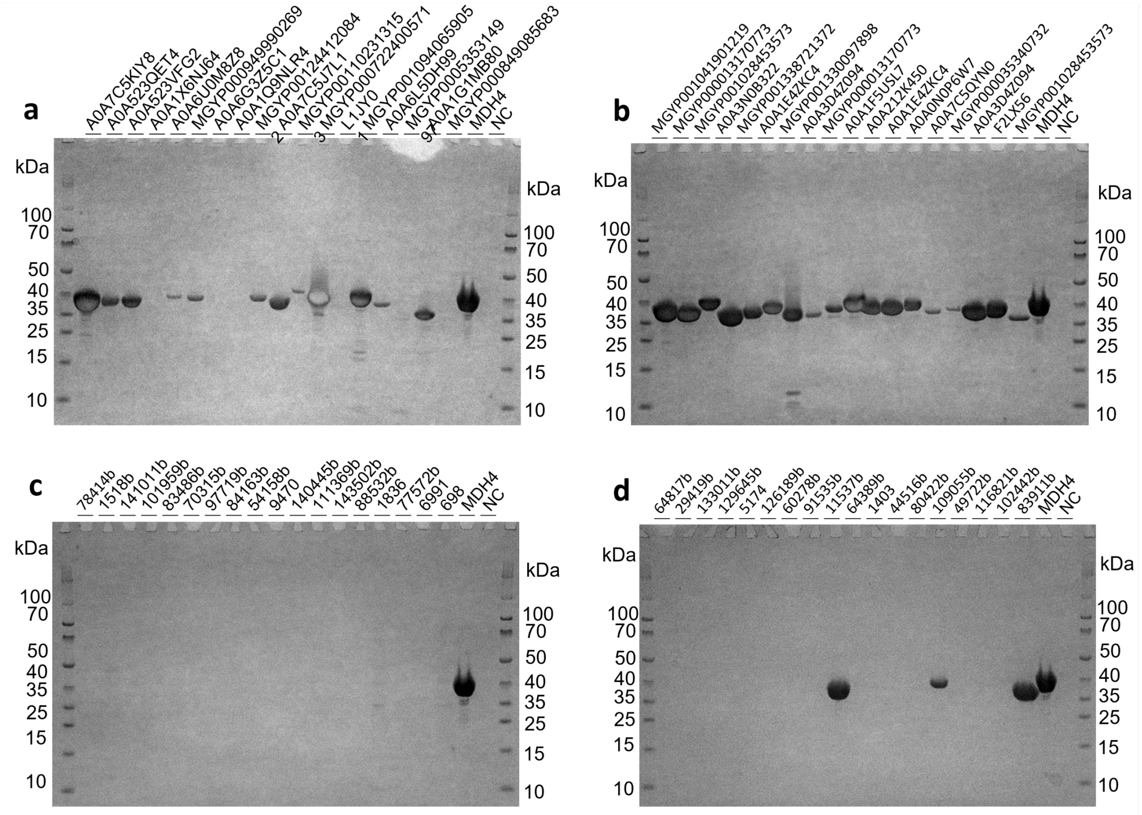


**Supplementary Fig 25. SDS gel of purified MDH enzymes in Round 3** from a) ESMMSA_control group, b) ESMMSA_selected group, c) GAN_control group and d) GAN_selected group.

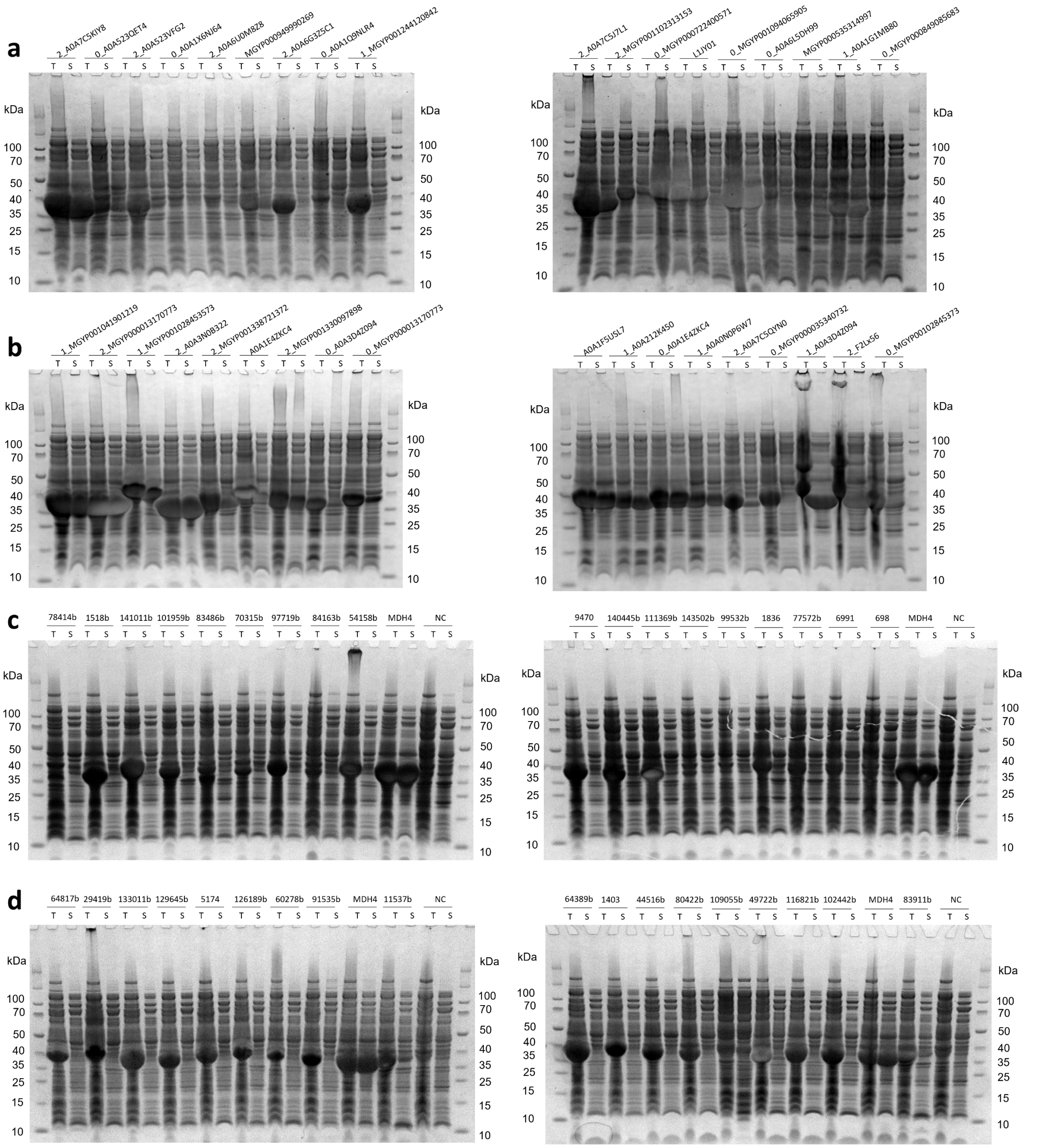

**Supplementary Fig 26. SDS gel of total (T) and soluble (S) MDH protein samples in Round 3** from a) ESMMSA_control group, b) ESMMSA_selected group, c) GAN_control group and d) GAN_selected group.

**a**

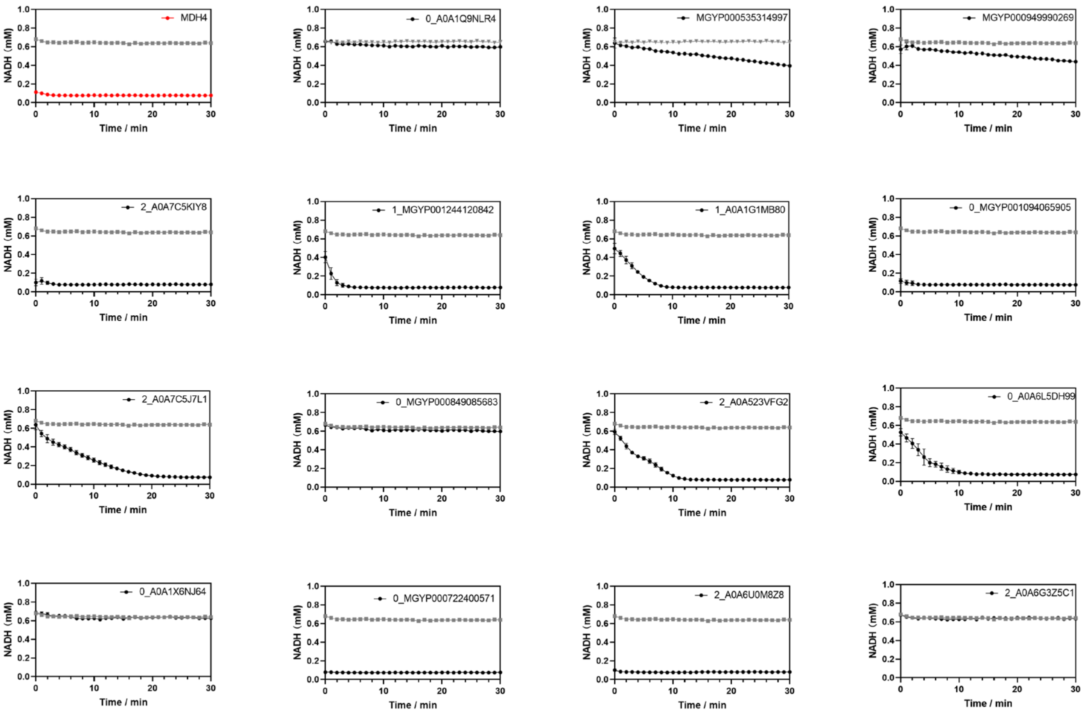
NADH (mM)
Time / min
MDH4
0_A0A1Q9NLR4
MGYP000535314997
MGYP000949990269
2_A0A7C5KIY8
1_MGYP001244120842
1_A0A1G1MB80
0_MGYP001094065905
2_A0A7C5J7L1
0_MGYP000849085683
2_A0A523VFG2
0_A0A6L5DH99
0_A0A1X6NJ64
0_MGYP000722400571
2_A0A6U0M8Z8
2_A0A6G3Z5C1

**b**

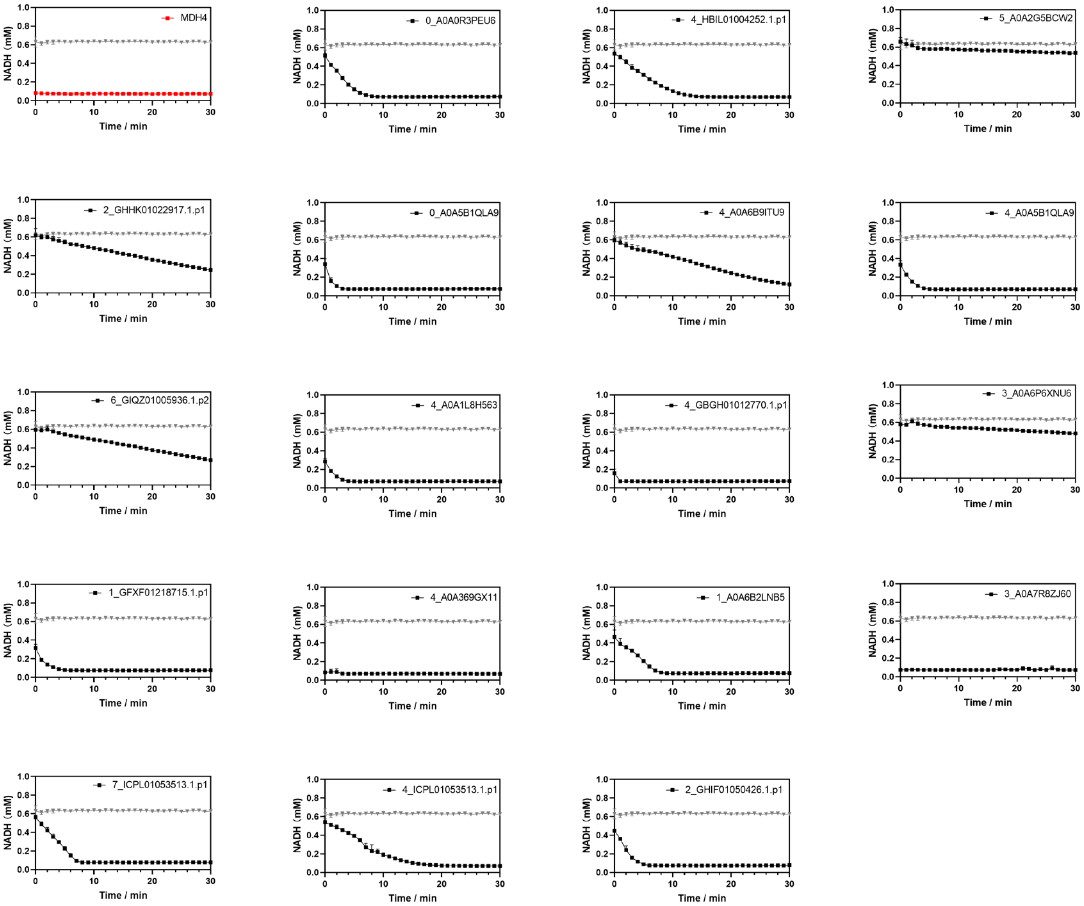
NADH (mM)
Time / min
MDH4
0_A0A0R3PEU6
4_HBIL01004252.1.p1
5_A0A2G5BCW2
2_GHHK01022917.1.p1
0_A0A5B1QLA9
4_A0A6B9ITU9
4_A0A5B1QLA9
6_GIQZ01005936.1.p2
4_A0A1L8H563
4_GBGH01012770.1.p1
3_A0A6P6XNU6
1_GFXF01218715.1.p1
4_A0A369GX11
1_A0A6B2LNB5
3_A0A7R8ZJ60
7_ICPL01053513.1.p1
4_ICPL01053513.1.p1
2_GHIF01050426.1.p1

**c**

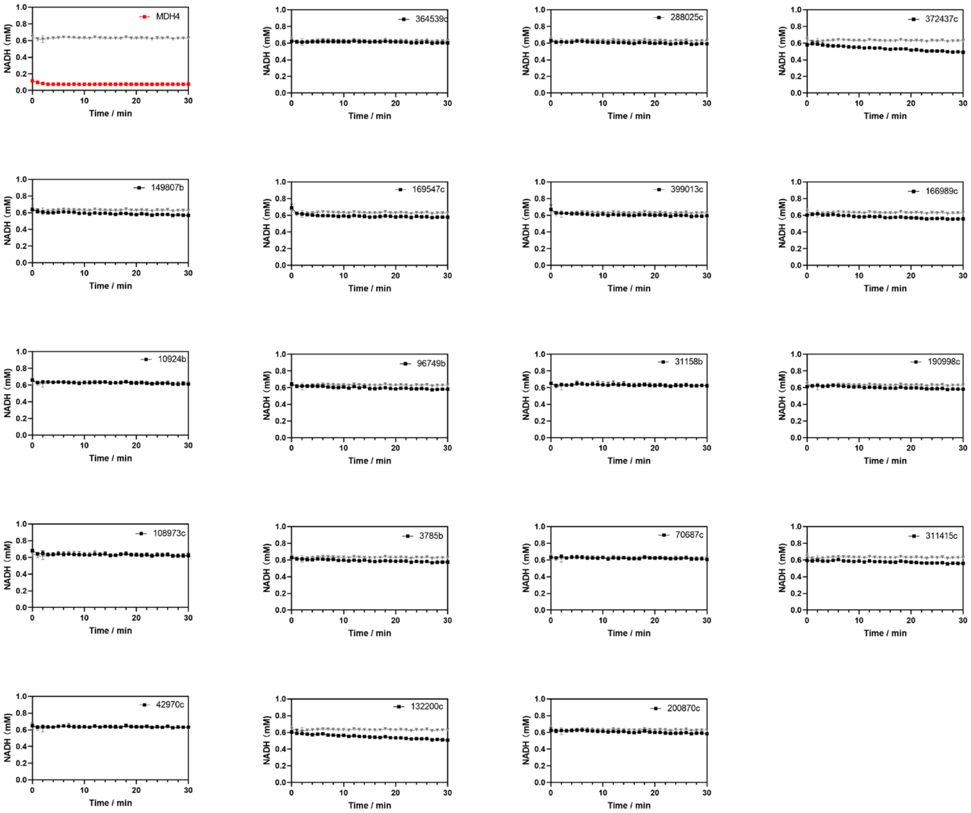
MDH4
364539c
288025c
372437c
149807b
169547c
399013c
166989c
10924b
96749b
31158b
190998c
108973c
3785b
70687c
311415c
42970c
132200c
200870c
NADH (mM)
Time / min

**d**

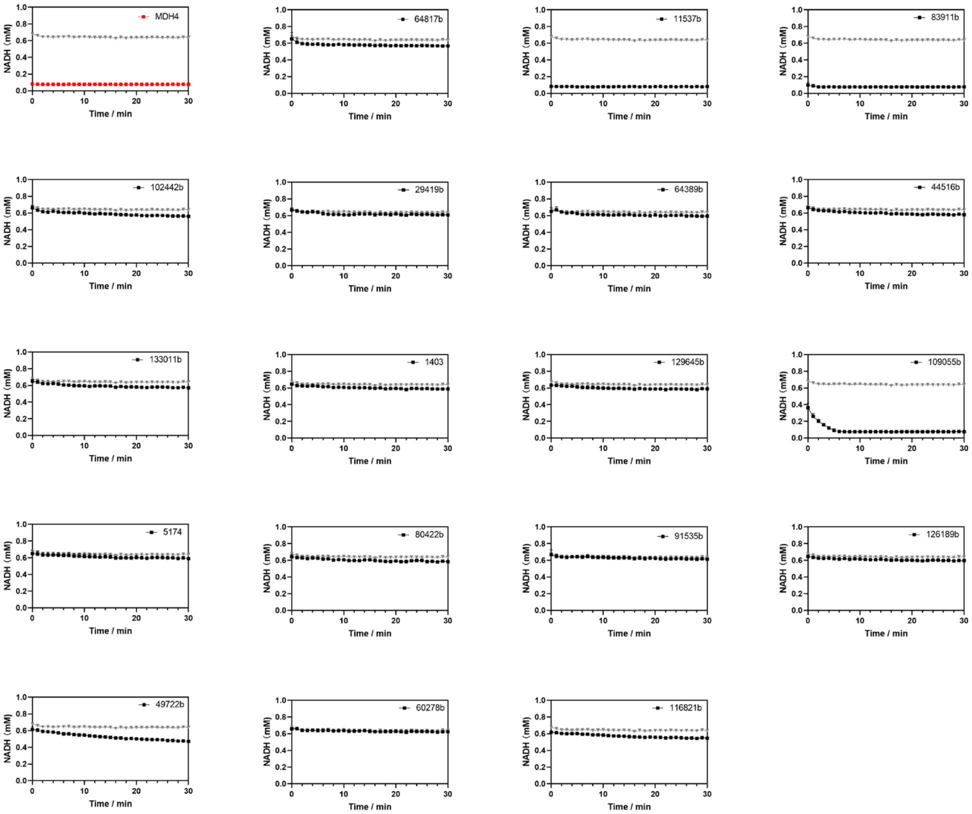
MDH4
64817b
11537b
83911b
102442b
29419b
64389b
44516b
133011b
1403
129645b
109055b
5174
80422b
91535b
126189b
49722b
60278b
116821b
NADH (mM)
Time / min

**Supplementary Fig 27. MDH enzyme activities in Round 3** from a) ESM-MSA control group, b) ESM-MSA selected group, c) GAN control group, d) GAN selected group measured in *vitro* by monitoring NADH consumption. An *E. coli* strain harboring pET-21b was used as a negative control. Wild-type malate dehydrogenases (MDH 4), likewise produced in *E. coli*. serves as positive controls. A time-dependent decrease in NADH can be readily observed for moderately active enzymes. For highly active enzymes all the NADH was consumed before measurements could take place, resulting in a horizontal line at the bottom of the plot. Negative controls are plotted in grey, positive controls in red, and generated sequences in black.

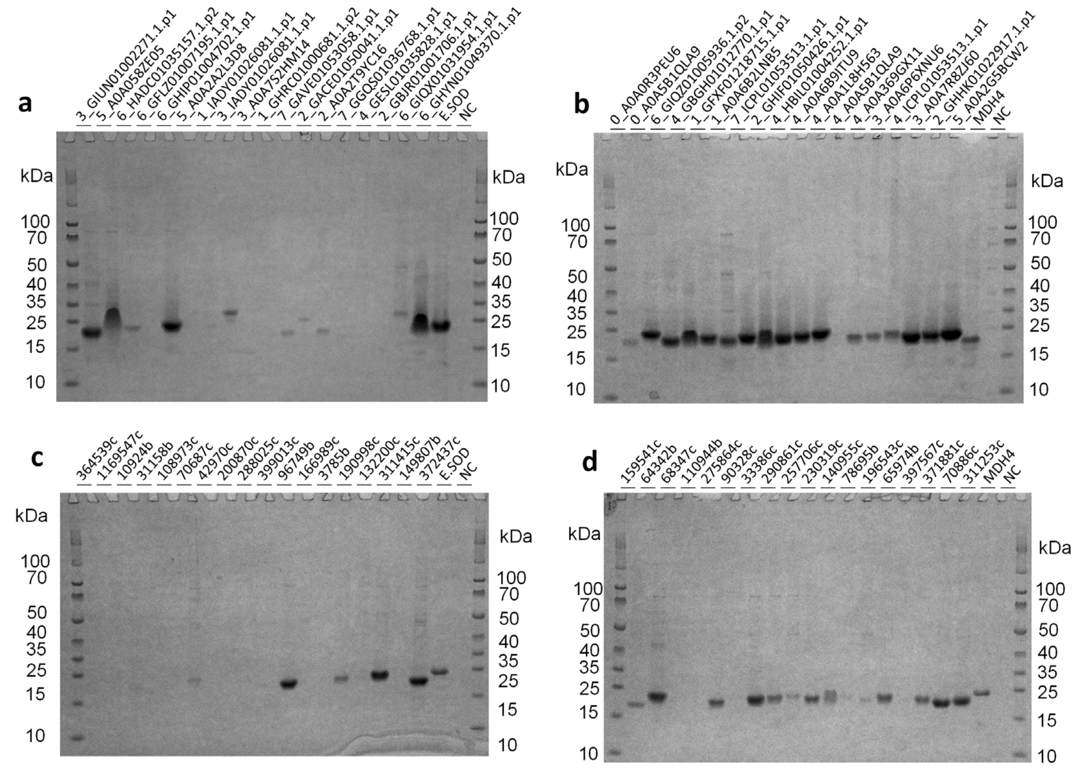


**Supplementary Fig 28. SDS gel of purified SOD enzymes in Round 3** from a) ESMMSA_control group, b) ESMMSA_selected group, c) GAN_control group and d) GAN_selected group.

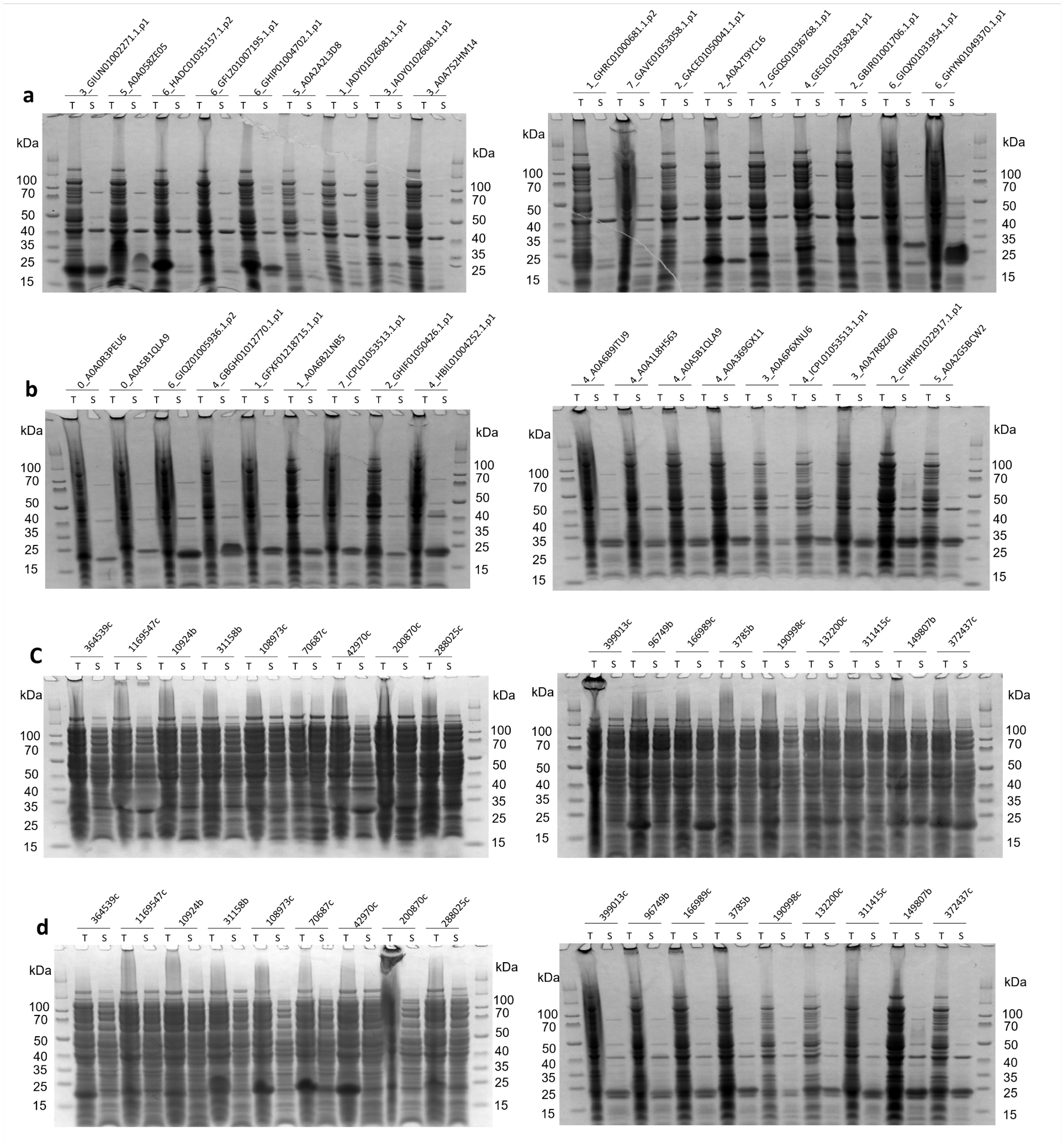


**Supplementary Fig 29. SDS gel of total (T) and soluble (S) SOD protein samples in Round 3** from a) ESMMSA_control group, b) ESMMSA_selected group, c) GAN_control group and d) GAN_selected group.

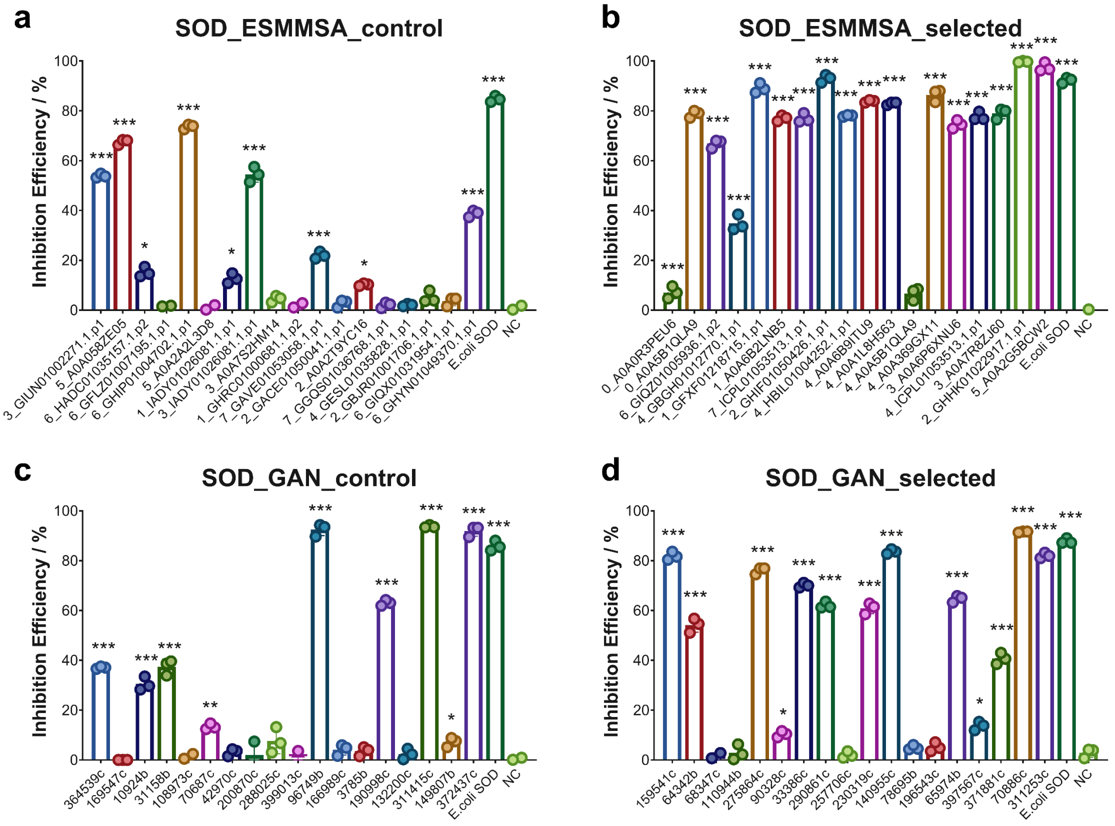


**Supplementary Fig 30. SOD enzyme activities in Round 3** from a) ESM-MSA control group, b) ESM-MSA selected group, c) GAN control group, d) GAN selected group, which were measured *in vitro* activity by the inhibition efficiency on xanthine oxidase. * marks samples with P-value less than 0.05 from the t-test compared to the negative control samples, ** for P < 0.01 and *** for P < 0.001.

### Supplementary Tables

#### Supplementary Table 1. Some possible failure modes for protein expression and activity.

| Category | Failure mode | Description |
|---|---|---|
| **Model failure**<br><br>These factors are particular to generated sequences and are less of a concern for natural sequences. | **Disruption of protein-protein binding residues** | For example, if a protein is active as part of a complex (either with identical or mixed subunits), mutations at the interface may cause misfolding or aggregation, or disrupt cooperative binding. |
| | **Mutation of catalytic residue to a non-catalytic residue** | Mutating a residue that participates in a reaction to a residue that can't participate in that reaction will cause loss of activity. |
| | **Mutations altering shape of active site or substrate/product/ligand binding sites or access tunnels** | Mutations to non-catalytic residues can disrupt enzyme function without affecting solubility/stability of a protein, such as by blocking a substrate or product channel or altering binding affinity to an important ligand. |
| | **Destabilizing mutation** | For example, mutating a buried position to a charged amino acid or to a large amino acid. |
| | **Aggregation-promoting mutation** | Hydrophobic patches on the protein surface, for example, can promote aggregation. |
| | **Mutation of cysteines involved in critical disulfide bridges** | The structures of some proteins are stabilized by disulfide bridges. Mutating the cysteines involved in disulfide bridges can destabilize a protein. Also keep in mind that *E. coli* cytosol is a poor environment for disulfide bridge formation, so proteins with critical disulfide bridges may fold poorly and be unstable in *E. coli* cytosol. |
| | **Disruption of sites for post translational modifications** | Some proteins depend on post-translational modifications, such as glycosylation or phosphorylation for folding or function. Mutating the modification sites can leave the mutants insoluble or inactive. |
| | **Disruption of protein dynamics** | Conformational changes are critical to the activity of many proteins and to the folding of all proteins. Mutations that disrupt protein dynamics, for example by locking a protein into a specific conformation, can disrupt function. |
| | **Insertions or deletions relative to functional reference proteins** | Deletions or insertions can dramatically change structure particularly if they occur in buried regions of the protein. |
| **Challenge of heterologous expression**<br><br>These factors can impact the expression and function of any protein, including natural proteins expressed outside of their native context. | **Lack of necessary chaperones, modifying proteins, binding proteins, redox partners, or ligands** | Some proteins require specific chaperones, enzymes to install ligands or post-translational modifications, proteins to form a heteromeric complex with, enzymes to regenerate redox state, or particular ligands. The necessary partners or ligands may be absent in *E. coli*. |
| | **Product toxicity** | Some proteins are toxic to *E. coli* and cannot be expressed in *E. coli* because they kill it. For example, nucleases and proteases can be difficult to express. |
| | **Presence of transmembrane domain** | Will cause the protein to stick in a membrane and therefore reduce amount of protein soluble in the cytosol. |
| | **Presence of signal peptides/localization tags not recognized or processed by *E. coli*.** | Many proteins have localization or secretion tags. In their native hosts, these tags are often cleaved off after the protein reaches its target organelle or compartment. Signal recognition machinery is specific to each lineage, so *E. coli* will fail to cleave most tags, leaving heterologous proteins as propeptides which could be inactive or unstable. |
| | **Long repeats** | Some generative models tend to produce long stretches of the same amino acid, amino acid pairs, or other low complexity patterns, which can lead to unstable proteins. |
| | **Sensitivity to pH, temperature, or ionic strength of expression media** | Some proteins, for example proteins secreted by extremophiles, may require chemical conditions very different from *E. coli* cytosol. |
| | **Issues with gene sequence** | The DNA sequence encoding a protein can also impact expression and solubility. For example, genes that consist mostly of codons that are rare in *E. coli* may express poorly. |

#### Supplementary Table 2. Curation of training data

| | CuSOD | | MDH | |
|---|---|---|---|---|
| | Round 1 | Rounds 2 and 3 | Round 1 | Rounds 2 and 3 |
| **Databases** | UniProt | UniProt, TSA Eukaryotes | UniProt | UniProt, Mgnify |
| **Phylogenetic origin** | no restriction | eukaryotic and viral | no restriction | no restriction |
| **Pfam domains** | Sod_Cu | Sod_Cu | Ldh_1_N and Ldh_1_C plus a better score against an HMM built from SwissProt MDH enzymes than an HMM built from LDH enzymes. | |
| **Sequence content** | no non-canonical amino acids | no non-canonical amino acids and starts with M | no non-canonical amino acids | no non-canonical amino acids and starts with M |
| **Phobius TM or SP** | | no predicted TM or SP | | no predicted TM or SP |
| **Truncation** | domain envelope plus up to 10 additional N-terminal amino acids, and 0 or 10 additional C-terminal amino acids. | | domain envelope plus up to 10 additional N-terminal amino acids, and 0 or 10 additional C-terminal amino acids. | |
| **Length filter** | throw out sequences > 1 standard deviation from the median. | throw out sequences > 1 standard deviation from the median, or extending more than 25 residues beyond the Sod_Cu envelope on the N-terminus, or 15 residues at the C-terminus. | throw out sequences > 1 standard deviation from the median. | throw out sequences > 1 standard deviation from the median, or extending more than 20 residues beyond the N-terminus of the Ldh_1_N domain envelope, or the C-terminus of the Ldh_1_C domain envelopes |
| **CD-HIT deduplication threshold** | 80% | 90% | 80% | 90% |
| **# of train sequences** | 4802 | 2543 | 3812 | 7187 |
| **# of test sequences** | 1201 | 283 | 953 | 799 |

#### Supplementary Table 3. Sequence generation

| | | Round 1 | | Rounds 2 and 3 | |
|---|---|---|---|---|---|
| | | CuSOD | MDH | CuSOD | MDH |
| **ASR** | Generated sequences | 443 | 883 | 945 | 973 |
| | MSA size | 445 | 835 | 947 | 975 |
| **GAN** | Generated sequences | 10048 | 10048 | 560064 | 160064 |
| | Note | | | 10048 sequences were generated for round 2, but additional sequences were generated for round 3. | |
| **ESM-MSA** | Generated sequences | 5000 | 5000 | 20344 | 28748 |
| | Sampling method | Random subset MSAs of 64 sequences randomly selected from the training MSA. Masking and sampling across the entire subset MSA. Two complete burn-in passes, and no top-k passes. | | MSAs of size 32, generated from top phmmer hits of each training sequence against the full list of training sequences. Mask and sample only the query sequence. Two burn-in passes followed by one top-k = 1 pass. | |
| | Note | | | One resampling of each training sequence (2543 CuSOD, and 7187 MDH) were generated for round2, but additional sequences were generated for round 3. | |

**Supplementary Table 4. Sequence selection** (* after the filters, the lowest identity selected sequence was 69% identical to the most similar training sequence).

| | **Percent identity to most similar training sequence** | **Metrics** |
|---|---|---|
| **Round 1** | 70-80 | Broad range of scores on ESM-1v unmasked metric. Manual inspection to remove sequences with large deletions or repeats. |
| **Round 2** | 80-90 | Broad range of scores on ESM-1v unmasked and ESM-MSA metrics. Manual inspection to remove sequences with large deletions or repeats. |
| **Round 3** | 50*-80 | Filter based on six criteria, including high scores from ESM-1v unmasked and ProteinMPNN. |

**Supplementary Table 5. Activity assay results**. Expressed: new visible band on SDS-PAGE gel of total protein compared with empty vector control. Soluble: new visible band on SDS-PAGE gel of soluble protein compared with empty vector control. Active: Measured activity significantly different from empty vector control. For some rows the total is less than 18, for the following reasons: in Round 1, three of the genes could not be synthesized by Twist; in Round 2, we selected only 13 CuSOD test sequences, because we had already tested some similar natural sequences in Round 2 pre-test.

| | Family | Model | Total | Expressed | Soluble | Active | Percent active |
|---|---|---|---|---|---|---|---|
| Round 1 | CuSOD | test | 17 | 5 | 3 | 0 | 0 |
| | | ASR | 18 | 12 | 9 | 9 | 50 |
| | | GAN | 18 | 10 | 5 | 2 | 11 |
| | | ESM-MSA | 18 | 12 | 4 | 0 | 0 |
| | MDH | test | 17 | 12 | 6 | 6 | 41 |
| | | ASR | 18 | 18 | 14 | 10 | 56 |
| | | GAN | 18 | 14 | 0 | 0 | 0 |
| | | ESM-MSA | 17 | 13 | 0 | 0 | 0 |
| Round 2 | CuSOD | test | 13 | 11 | 11 | 7 | 54 |
| | | ASR | 18 | 18 | 18 | 15 | 83 |
| | | GAN | 18 | 18 | 18 | 12 | 67 |
| | | ESM-MSA | 18 | 18 | 18 | 12 | 67 |
| | MDH | test | 18 | 18 | 15 | 14 | 78 |
| | | ASR | 18 | 16 | 16 | 13 | 72 |
| | | GAN | 18 | 15 | 3 | 2 | 11 |
| | | ESM-MSA | 18 | 17 | 12 | 9 | 50 |
| Round 3 | CuSOD | GAN-passing | 18 | 16 | 16 | 13 | 72 |
| | | GAN-control | 18 | 11 | 9 | 9 | 50 |
| | | ESM-MSA-passing | 18 | 18 | 18 | 17 | 94 |
| | | ESM-MSA-control | 18 | 11 | 11 | 9 | 50 |
| | MDH | GAN-passing | 18 | 4 | 4 | 4 | 22 |
| | | GAN-control | 18 | 0 | 0 | 2 | 11 |
| | | ESM-MSA-passing | 18 | 18 | 18 | 18 | 100 |
| | | ESM-MSA-control | 18 | 14 | 12 | 12 | 67 |

#### Supplementary Table 6. Details of COMPSS validation on six additional datasets

| | Chorismate mutase | Phage lysozyme | Glucosaminidase | Glycosidase hydrolase | Transglycosylase | Pesticin |
|---|---|---|---|---|---|---|
| **Pfam accession** | PF01817 | PF00959 | PF01832 | PF05838 | PF06737 | PF16754 |
| **Training set curation** | From reference | Hmmsearch of Pfam profile against UniProt, extraction of the domain plus padding on each side, clustering at 90% identity. | | | | |
| **Number of training sequences** | 1130 | 8023 | 16568 | 3514 | 5800 | 830 |
| **ESM-1v top 10% threshold** | -0.28 | -0.24 | -0.19 | -0.20 | -0.11 | -0.36 |
| **Generative model** | bmDCA | ProGen | | | | |
| **Reference** | Russ et al., 2020 | Madani et al., 2023 | | | | |
| **ESM-1v AUC-ROC** | 0.75 | 0.30 | 0.20 | 0.60 | 1.00 | 0.60 |
| **ProteinMPNN AUC-ROC** | 0.77 | 0.63 | 1.00 | 0.60 | 0.78 | 0.81 |
| **COMPSS filter** | ESM-1v score in top 10% of training set, no predicted transmembrane domain, longest single repeat shorter than 4, longest pair repeat shorter than 6. We skipped the 'starts with M' filter because very few of the sequences in these sets start with M, and did not subset by identity to closest training sequence. | | | | | |
| **COMPSS passing, active/total (% active)** | 82 / 130 (63%) | 1 / 2 (50%) | 1 / 1 (100%) | 2 / 2 (100%) | 1 / 1 (100%) | 1 / 1 (100%) |
| **COMPSS failing, active/total (% active)** | 400 / 1487 (27%) | 14 / 17 (82%) | 4 / 5 (80%) | 32 / 41 (78%) | 2 / 5 (40%) | 8 / 15 (53%) |

#### Supplementary Table 7. List of PCR primers

| primer name | sequence |
|---|---|
| hSOD_F | AAGAAGGAGATATACATATGGCGACGAAGGCCGTG |
| hSOD_R | TGCTCGAGTGCGGCCGCTTGGGCGATCCCAATTAC |
| hSOD_Ntrun_F | AAGGAGATATACATATG GCCGTGTGCGTGCTGAAG |
| hSOD_Ctrun_R | TGCTCGAGTGCGGCCGCAATTACACCACAAGCCAA |
| paSOD_F | AAGAAGGAGATATACATATGGCGAAAGGCGTTGCT |
| paSOD_R | GTGCTCGAGTGCGGCCGCTCCCTGCAACCCGATGA |
| paSOD_Ntrun_F | AAGGAGATATACATATGGCGTTGCTGTGCTTTCTT |
| paSOD_Ctrun_R | TGCTCGAGTGCGGCCGCGATGATTCCACATGCGAT |
| E.coliSOD_F | AAGGAGATATACATATGTCATTCGAATTACCTGCA |
| E.coliSOD_Ctrun_R | TGCTCGAGTGCGGCCGCATTTTTCGCTACGAATTC |
| T7_forward | TAATACGACTCACTATAGGG |
| T7term_reverse | CTAGTTATTGCTCAGCGGT |
